# Intracellular nanovesicles mediate integrin trafficking during cell migration

**DOI:** 10.1101/2020.08.19.257287

**Authors:** Gabrielle Larocque, Penelope J. La-Borde, Beverley J. Wilson, Nicholas I. Clarke, Daniel J. Moore, Patrick T. Caswell, Stephen J. Royle

## Abstract

Membrane traffic is an important regulator of cell migration through the endocytosis and recycling of cell surface receptors such as integrin heterodimers. Intracellular nanovesicles (INVs), are a recently identified class of transport vesicle that are involved in multiple membrane trafficking steps including the recycling pathway. The only known marker for INVs is Tumor Protein D54 (TPD54/TPD52L2), a member of the TPD52-like protein family. Overexpression of TPD52-like family proteins in cancer has been linked to poor prognosis and an aggressive metastatic phenotype which suggests cell migration may be altered under these conditions. Here we show that TPD54 associates with INVs by directly binding high curvature membrane via a conserved positively charged motif in its C-terminus. We describe how other members of the TPD52-like family are also associated with INVs and we document the Rab GTPase complement of all INVs. Depletion of TPD52-like proteins inhibits cell migration and invasion; and we show that this is likely due to altered integrin recycling. Our study highlights the involvement of INVs in the trafficking of cell surface proteins to generate biologically important outputs in health and disease.

## Introduction

Cell migration is important for many aspects of animal physiology: the immune response, tissue integrity and embryonic development. This process is tightly controlled and any alterations can result in diseases such as inflammation or cancer (Hamidi and Ivaska, 2018; Friedl and Wolf, 2003). Membrane traffic is a key regulator of cell migration through the trafficking of cell surface receptors, including integrins, which bind the extracellular matrix (ECM). Integrins are endocytosed and recycled back to the cell surface in order to break and re-establish the cellular contacts with the ECM during migration. The molecular details of integrin trafficking pathways and their influence on cell motility are under active investigation (Wilson et al., 2018).

The identity and activation state of integrin heterodimers govern their trafficking and fate (Wilson et al., 2018). Rabs are master regulators of membrane traffic with >60 different Rabs in human cells, with each one mediating a trafficking step specifically (Wandinger-Ness and Zerial, 2014). Rabs are involved in every step of integrin traffic, as well as binding integrins directly or via one of the many Rab effector proteins (Pellinen et al., 2006; Caswell et al., 2008).

A new class of intracellular transport vesicle, termed intracellular nanovesicles (INVs), has recently been described (Larocque et al., 2020). INVs are involved in recycling and anterograde trafficking pathways. These vesicles are small (~30nm diameter), highly dynamic and are associated collectively with ~16 different Rab GTPases (Larocque et al., 2020). Among the Rabs present on INVs are Rab11a and Rab25, two Rabs well known for their involvement in integrin trafficking (Roberts et al., 2001; Caswell et al., 2007; Moreno-Layseca et al., 2019). INVs were discovered because of their association with TPD54, a member of the Tumor Protein D52-like protein family (TPD52, TPD53/TPD52L1, TPD54/TPD52L2 and TPD55/TPD52L3). How TPD54 associates with INVs and whether the other members of the TPD52-like protein family behave similarly are important open questions.

TPD52-like proteins were identified due to their overexpression in a number of cancer types (Byrne et al., 1995, 1996; Nourse et al., 1998; Cao et al., 2006). This upregulation is often caused by gene duplication which is tumorigenic (Balleine et al., 2000; Lewis et al., 2007). Tumorigenicity has been proposed to be due to alteration of either the cell cycle (Boutros and Byrne, 2005; Thomas et al., 2010; Lewis et al., 2007), signaling (Li et al., 2017), or DNA repair (Chen et al., 2013). Finally, TPD52-like proteins have been reported to have a role in cell migration and adhesion, however, the underlying mechanism is unknown (Ummanni et al., 2008; Mukudai et al., 2013). In breast cancer, TPD52 overexpression correlates with poor prognosis a decrease in metastasis-free survival (Roslan et al., 2014; Shehata et al., 2008). This suggested to us that TPD52-like proteins, and the INVs they are associated with, may be involved in cell migration and invasion in cancer.

In this paper we show how TPD52-like proteins associate with INVs and document the Rab complement of INVs decorated with TPD52, TPD53 or TPD54. We find that depletion of TPD52-like proteins inhibits cell migration and invasion, and show that this is likely due to altered integrin recycling via INVs.

## Results

### Molecular determinants required for the association of TPD54 with INVs

We previously found that TPD54 is tightly associated with INVs and that its association could be measured by spatiotemporal variance of fluorescence microscopy images (Larocque et al., 2020). Here, we asked what are the molecular determinants for the association of TPD54 with INVs. Analysis of the primary sequences of TPD52-like proteins reveals two domains. First, a coiled-coil domain between residues 38 and 82 (Figure 1A). Second, a region between residues 126 and 180 with high similarity among human TPD52-like proteins. Within this region, residues 160-171 (Figure 1A) are particularly well conserved across different species (Supplementary Figure S1). With these regions in mind, we designed mCherry-FKBP-tagged TPD54 constructs to pinpoint which region of the protein was required for its association with INVs (Figure 1B). The spatiotemporal variance of fluorescence for mCherry-FKBP-TPD54 full-length was much higher than that of the control mCherry-FKBP, which is to be expected due to the association of TPD54 with mobile, sub-resolution vesicles such as INVs (Figure 1B and Supplementary Video SV1). The variance of the TPD54 constructs revealed that the C-terminal portion of TPD54 (155-180) was needed for INV localization (Figure 1B). Constructs that lacked this region (1-37, 1-82, 38-82, 1-155) had low variance, while those that contained it (1-206, 83-206, 1-180) had higher variance.

**Figure 1.**
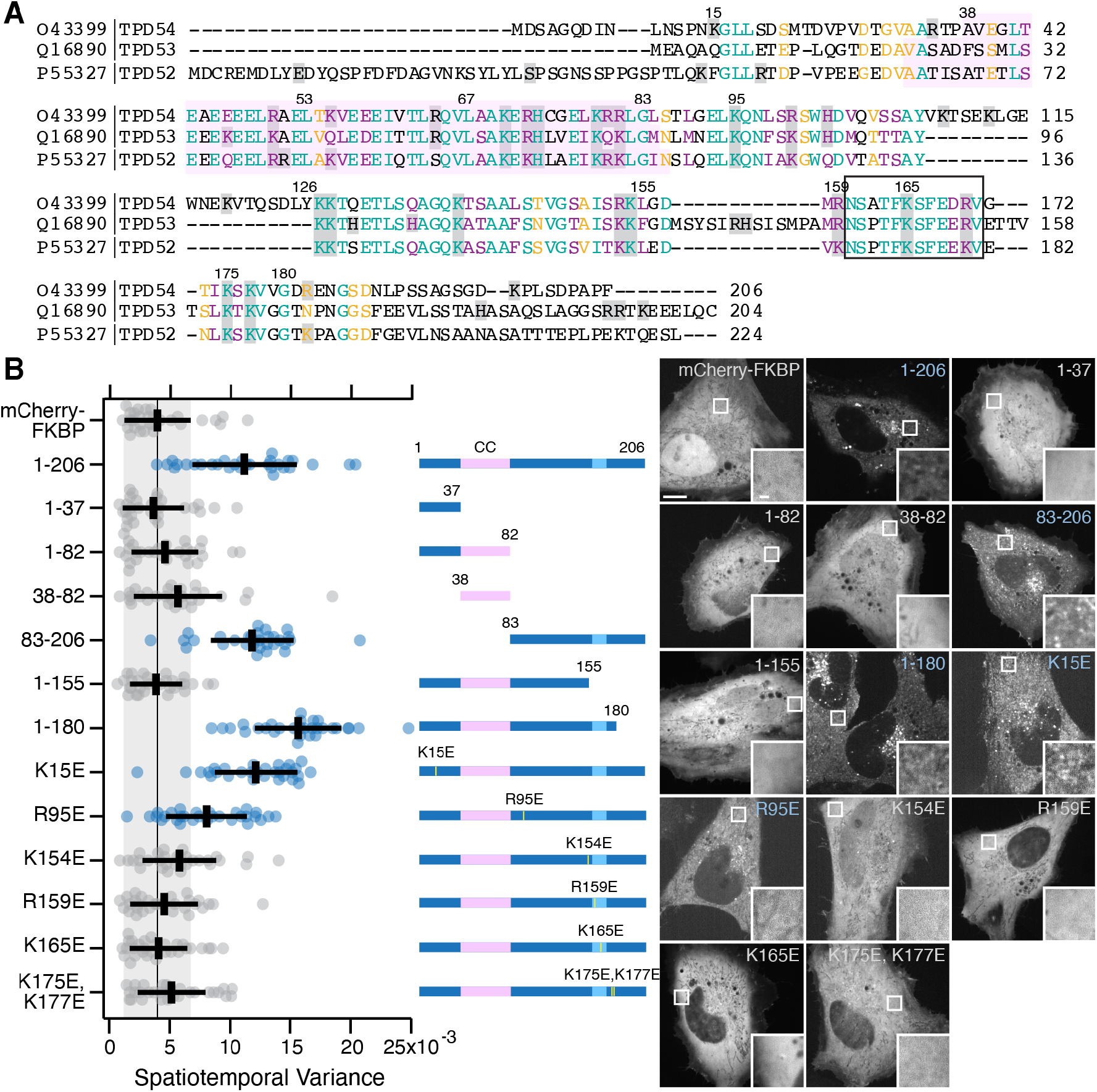
Identification of the region required for the association of TPD54 with INVs. (**A**) Alignment of human TPD54/TPD52L2, TPD53/TPD52L1 and TDP52/TPD52. Numbers indicate residue position in TPD54. Pink highlighted area, coiled-coil domain. Boxed area, conserved region (see Supplementary Figure S1). Gray shadowing shows positively charged residues. Lettering: Teal, fully conserved residues; Purple, strongly similar (>0.5 in the Gonnet PAM 250 matrix); Yellow, weakly similar (<0.5 in the Gonnet PAM 250 matrix). (**B**) Scatter dot plot to show the mean variance per pixel over time for the indicated constructs. Dots, individual cells; Black bars, mean ±sd. The mean ±sd for mCherry-FKBP (control) is also shown as a black line and gray zone, down the plot. Dunnett’s *post-hoc* test was done using mCherry-FKBP as control; blue indicates *p* < 0.05. A schematic representation of the mCherry-FKBP-tagged TPD54 constructs used in this figure. Pink region, coiled-coil domain (CC). Yellow line, position of point mutation. Light blue region, boxed area in (A). Representative confocal micrographs of HeLa cells expressing mCherry-FKBP or mCherry-FKBP-tagged TPD54 FL (1-206) or indicated mutants. Blue labels indicate high variance measured in B. Inset: 4× zoom. Scale bar, 10μm or 1 μm (inset).

The region around residues 155-180 contains several positively charged residues (Figure 1A), suggesting that this region could bind direct to the INV membrane. Mutation of either K154, R159, K165 and K175/K177 to glutamic acid reduced the variance to levels similar to control (Figure 1B). Whereas similar mutations (K15E or R95E) outside this region had no significant effect. These experiments establish that positively charged residues around the conserved 155-180 region of TPD54 are responsible for its association with INVs.

### TPD54 directly and preferentially binds high curvature vesicles

The association of TPD54 with INVs could either be direct or via another membrane associated protein. We therefore investigated whether or not TPD54 was able to bind vesicles directly, using a liposome-binding assay. From a Folch extract of brain lipids, we generated liposomes of four different sizes, 30, 50 100 or 200 nm and tested for co-sedimentation of GST, GST-TPD54 or GST-TPD54(R159E) with these liposomes. We observed specific co-sedimentation of both TPD54 proteins, but not GST alone, with liposomes of all sizes (Figure 2A). In order to quantify this direct membrane binding, we first measured the efficiency of liposome pelleting since smaller liposomes sediment less efficiently (Boucrot et al., 2012) and are less abundant (see Methods, Figure 2B,C). Quantification of co-sedimentation, accounting for liposome pelleting efficiency, revealed that GST-TPD54 bound 30 nm and 50 nm tighter than to larger diameter liposomes (Figure 2A,D). The binding of GST-TPD54(R159E) to all sizes of liposomes was lower than that observed for WT. Moreover, the R159E mutant had no size preference for liposomes of high curvature (Figure 2A,D).

**Figure 2.**
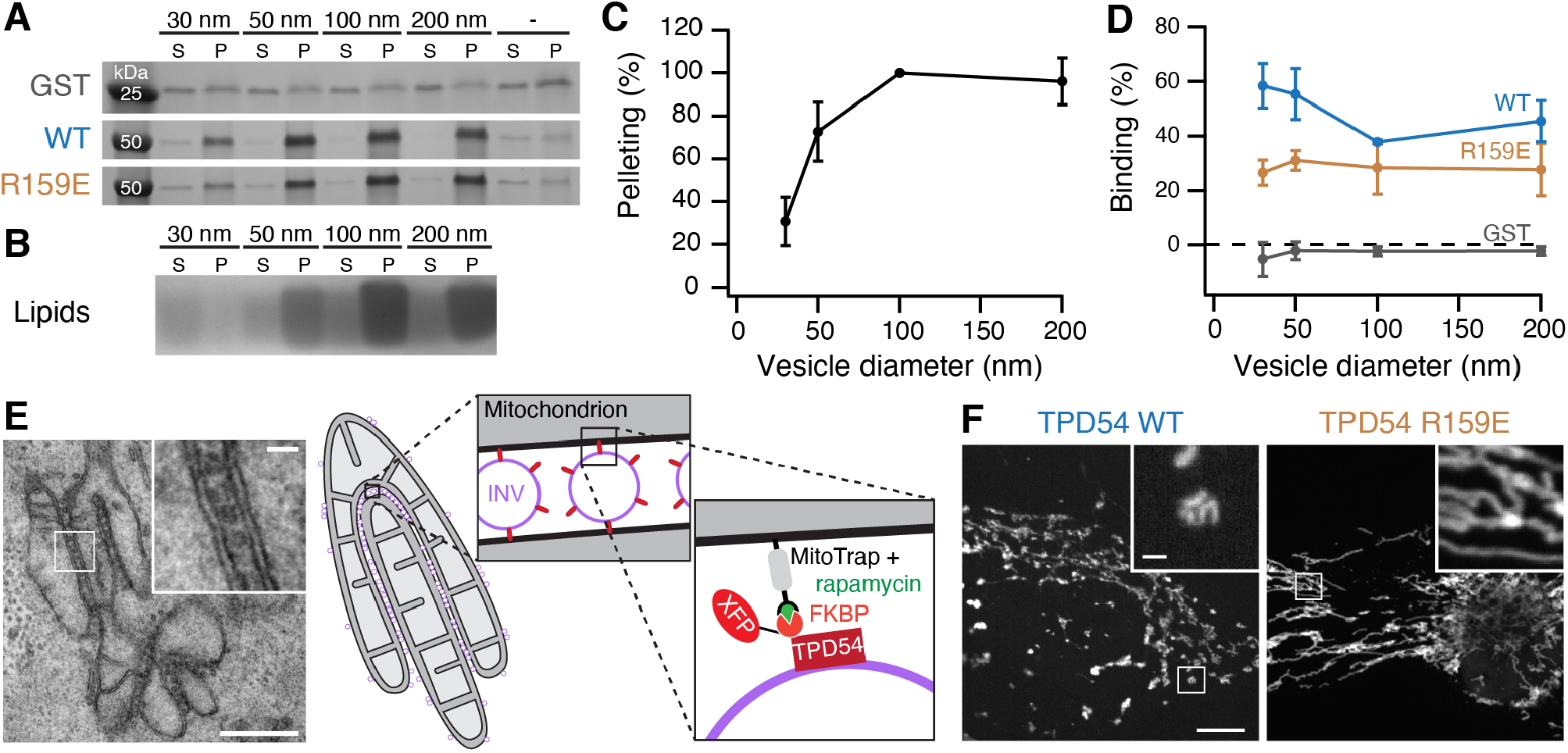
TPD54 binds directly to INVs. (**A**) Co-sedimentation of GST, GST-TPD54 WT, or GST-TPD54(R159E) with differently sized liposomes (diameter indicated) or no liposomes (-), visualized on an InstantBlue-stained 4% to 12% gel. (**B**) Pelleting of differently sized liposomes. Coomassie-stained Bis-Tris gel showing lipid stain behind the dye front. S, supernatant; P, pellet. (**C**) Quantification of the pelleting efficiency of liposomes according to their size. Values are normalized to 100 nm liposomes. (**D**) Quantification of protein co-sedimentation with differently sized liposomes. The binding of GST (gray), GST-TPD54 (WT, blue) or GST-TPD54(R159E, gold) is shown corrected for the pelleting efficiency of liposomes and for the sedimentation of the proteins without liposomes. Bars, s.d. *n*_exp_ = 3. (**E**) Capture of INVs causes mitochondrial aggregation. Electron micrograph of a cell where GFP-FKBP-TPD54 WT was rerouted to MitoTrap with schematic representation (right). Inset: 4× zoom. Scale bar, 500 nm; inset, 50 nm. An expanded version of this panel is shown in Supplementary Figure S2. (**F**) Representative confocal micrographs of HeLa cells expressing dark MitoTrap and either mCherry-FKBP-TPD54 WT or R159E mutant, treated with 200 nM rapamycin. Inset: 5× zoom. Scale bar, 10 μm, 1 μm (inset).

In order to verify that the R159E mutation prevented association of TPD54 with INVs in cells, we used a simple visual assay (Figure 2E,F). Previously, we found that rerouting an FKBP-tagged TPD54 construct to MitoTrap on mitochondria using rapamycin, causes mitochondrial aggregation due to capture of INVs (Larocque et al., 2020). Rerouting mCherry-FKBP-TPD54 to MitoTrap in HeLa cells caused mitochondrial aggregation; whereas the mitochondria were unaffected by rerouting the mCherry-FKBP-TPD54(R159E) (Figure 2F). These results indicate that TPD54 binds membrane directly, and that it has a preference for high curvature vesicles. The association of TPD54 with INVs in cells is likely via an interaction between negatively charged lipids and the positive residues surrounding the conserved 155-180 region.

### TPD52-like proteins can homo- and heteromerize via the coiled-coil domain, but multimerization is dispensable for INV binding

It is likely that TPD54 can form homomers or heteromers with other TPD52-like proteins via their coiled-coil domains (Byrne et al., 1998; Sathasivam et al., 2001; Larocque et al., 2020). This raises the question of whether the association with INVs requires multimerization. In order to answer this point, we sought a TPD54 mutant that was incapable of multimerization. Mutation of two leucines to prolines is predicted to break the coiled-coil domain of TPD54 (L53P,L67P Figure 1A and 3A) and so we tested whether these mutations interfered with homo- and heteromerization in cells. FLAG-tagged TPD52-like proteins were immunoprecipitated from cells co-expressing either GFP-FKBP, GFP-FKBP-TPD54 or GFP-FKBP-TPD54(L53P,L67P). We found that TPD54 WT, but neither the control nor the mutant could be co-immunoprecipitated with FLAG-tagged TPD52, TPD53 or TPD54 (Figure 3B). These results confirmed that the coiled-coil domain of TPD54 is responsible for homo- and heteromerization and that the L53P,L67P mutation blocks multimerization. Next, we found that the localization of the L53P,L67P mutant was normal and its spatiotemporal variance was similar to wild-type TPD54, suggesting that it was associated with INVs as normal (Figure 3C,D). This result indicates that monomeric TPD54 can associate with INVs and that multimerization is not required for membrane binding.

**Figure 3.**
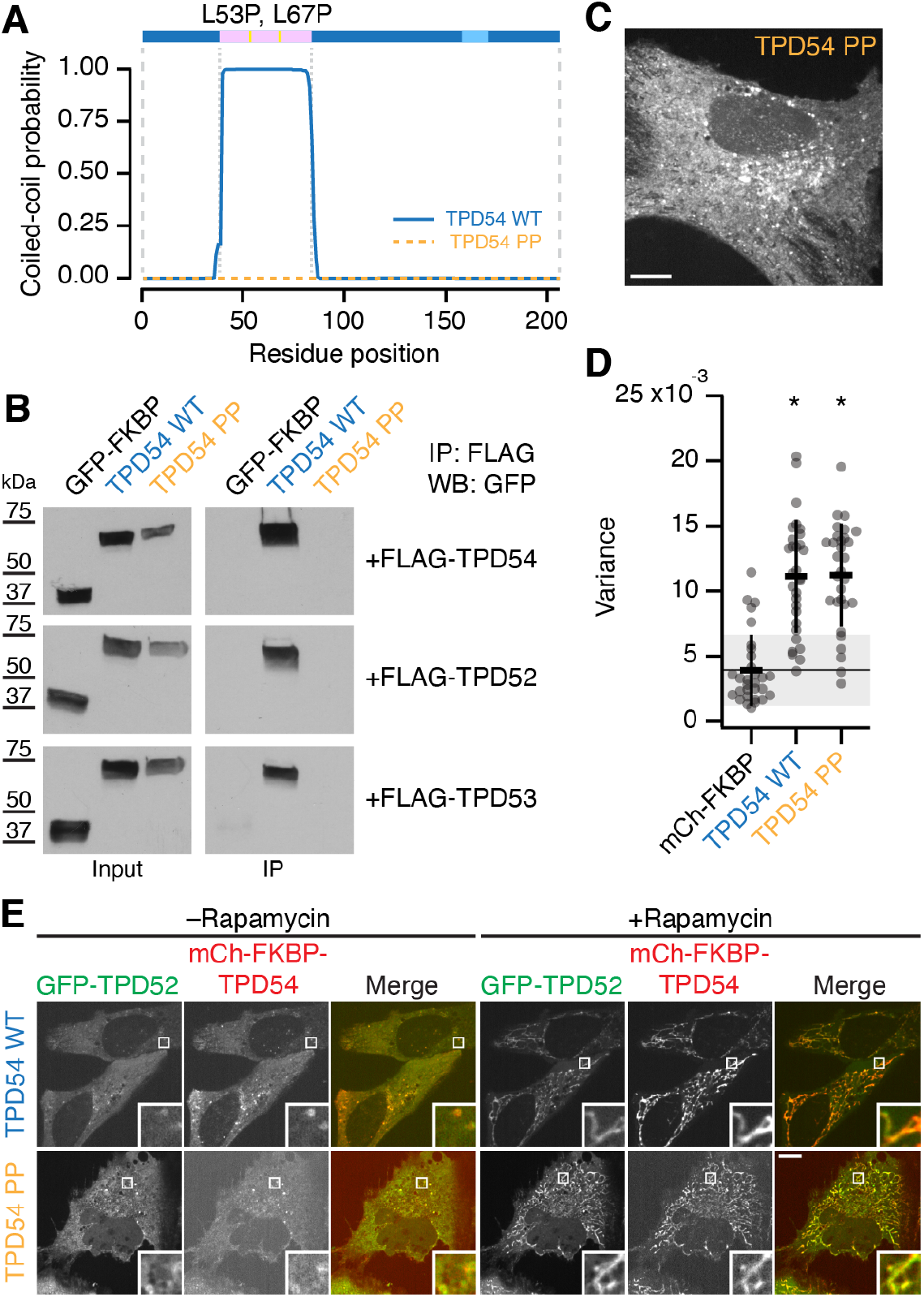
TPD54 homo- and heteromerizes using its coiled-coil domain. (**A**) Schematic representation of TPD54 and a graph showing the coiled-coil probability for wild-type (WT) and L53P,L67P (PP) mutant. Pink, coiled-coil domain; Light blue, conserved region between residues 158-171. For coiled-coil domain prediction, the amino acid sequence of TPD54 isoform 1 WT or TPD54 L53P, L67P was analyzed by PCOILS with window size of 28 (Gruber et al., 2006). (**B**) Western blot (WB) showing immunoprecipitation (IP) of FLAG-TPDs and co-immunoprecipitation of GFP-FKBP, GFP-FKBP-TPD54 (WT) or GFP-FKBP-TPD54(L53P,L67P) (PP). (**C**) Representative confocal image of mCherry-FKBP-TPD54(L53P,L67P) (PP)expressed in HeLa cell. Scale bar, 10 μm. (**D**) Scatter dot plot to show the mean variance per pixel over time. Dots, individual cells; black bars, mean ±sd. The mean ±sd for mCherry-FKBP (control) is also shown as a black line and gray zone. *, *p* < 0.05; Dunnett’s post-hoc test using mCherry-FKBP as control. Note, this is part of the dataset shown in Figure 1. (**E**) Representative confocal micrographs showing the co-rerouting of GFP-TPD52 after rerouting of mCherry-FKBP-TPD54 (top) or mCherry-FKBP-TPD54(L53P,L67P) (PP) (bottom) to dark Mitotrap by addition of 200 nM rapamycin. Inset: 5× zoom. Scale bar, 10 μm.

Given the similarity of TPD52-like proteins and the conservation of the residues involved in membrane association (Figure 1), it is likely that TPD52 and TPD53 are also found on INVs. Indeed, live cell imaging indicates that they have a similar subcellular distribution and spatiotemporal variance (Supplementary Video SV2). The TPD54(L53P,L67P) mutant allowed us to ask whether other TPD52-like proteins are associated with INVs independently of TPD54. To do this we used the mitochondrial INV-capture procedure using mCherry-FKBP-TPD54 or mCherry-FKBP-TPD54(L53P,L67P). We found that when GFP-TPD52 is co-expressed, it also becomes rerouted to the mitochondria (Figure 3E). This indicates that not only is TPD52 on INVs, but its association is likely direct and not via recruitment by TPD54.

### TPD52 and TPD53 are associated with a different subset of INVs

INVs are involved in many trafficking pathways since collectively they have a variety of Rab GTPases (Larocque et al., 2020). This was demonstrated by using mCherry-FKBP-TPD54 in a vesicle capture assay and asking which GFP-Rabs were co-rerouted to mitochondria. Since TPD52 and TPD53 also are on INVs (Figure 3E, Supplementary Video SV2 and Supplementary Figure S2), we wanted to know if all INVs have TPD54 or if there are subsets of INVs with different TPD52-like proteins. To investigate this point, we carried out mitochondrial vesicle capture and quantified the co-rerouting of GFP-Rabs to mitochondria using either mCherry-FKBP-TPD52 (Figure 4A,B) or mCherry-FKBP-TPD53 (Figure 4C,D). Of the 39 Rabs tested, 16 co-rerouted with TPD52 and 9 co-rerouted with TPD53. As with TPD54, Rab30 was the strongest hit with TPD52 (Figure 4B) and TPD53 4D), this suggests that Rab30 is the Rab GTPase most likely to be associated with INVs and can be thought of as an “INV Rab”. The other Rabs co-rerouted with TPD52 were Rab14, Rab26, Rab1a, Rab1b, Rab10, Rab17, Rab33a, Rab19b, Rab4a, Rab3a, Rab25, Rab21, Rab12 and Rab43. Of these, only Rab21 had not been identified in the TPD54 screen (Larocque et al., 2020). The Rabs co-rerouted with TPD53 were Rab30, Rab1b, Rab26, Rab1a, Rab33b, Rab43, Rab19b, Rab14, Rab12, and Rab10; all of which also co-rerouted with both TPD54 and TPD52. To classify INVs in a more stringent manner, we used hierarchical clustering of the mean mitochondrial intensity change of GFP-Rabs in the TPD52 and TPD53 vesicle capture screens (Figure 4A,C), together with the previously published TPD54 screen (Figure 5A). This resulted in the classification shown in Figure 5B.

**Figure 4.**
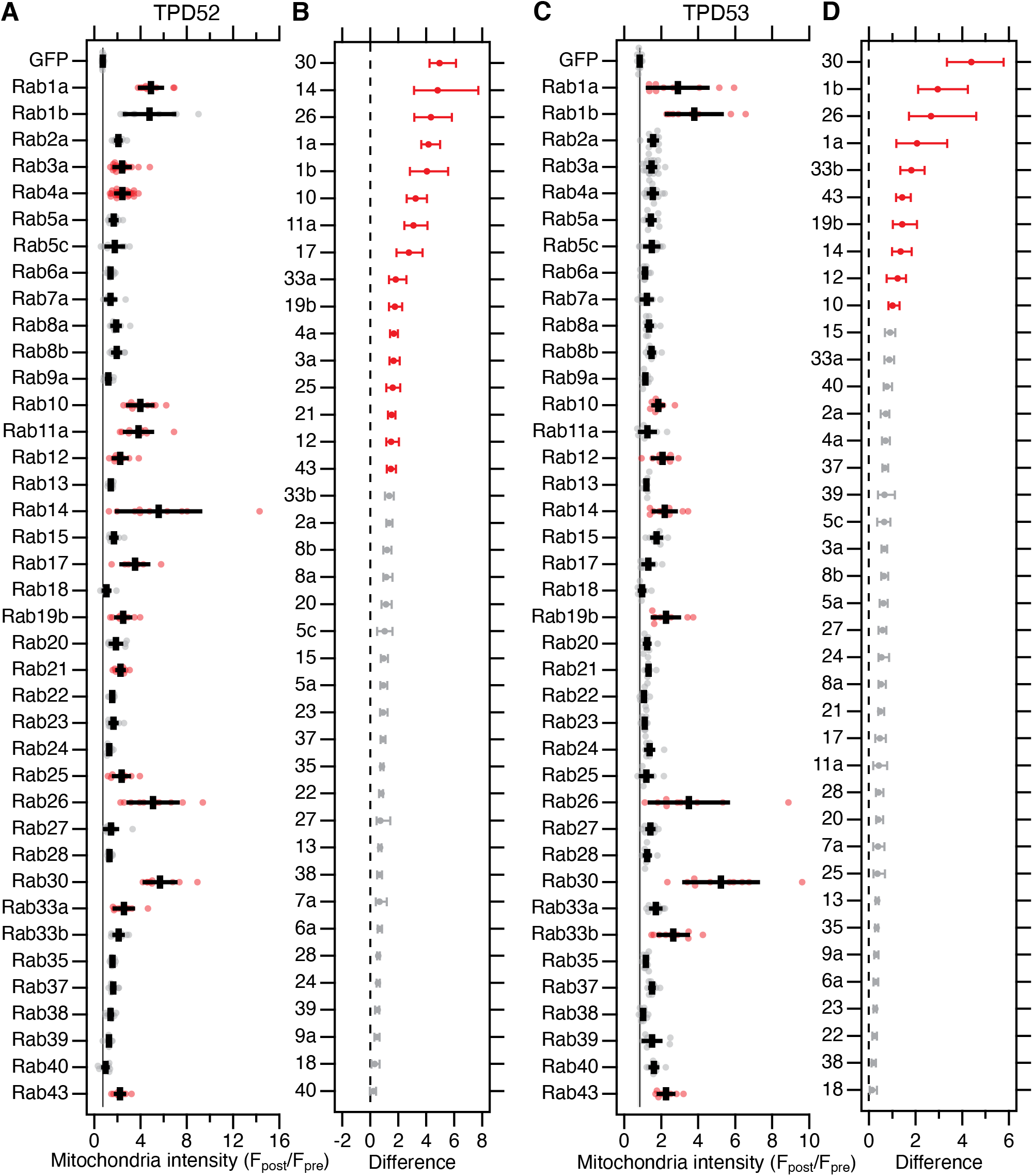
Screening Rab GTPases that are associated with TPD52 and TPD53 INVs. (**A,C**) Quantification of the change in mitochondrial fluorescence intensity of GFP or GFP-Rabs 2min after rerouting of mCherry-FKBP-TPD52 (A) or mCherry-FKBP-TPD53 (C) to dark MitoTrap with 200nM rapamycin. Dots, values for individual cells across three independent trials. Black bars, mean ±sd. The mean ±sd forGFP (control) is also shown as a black line and gray zone, down the plot. Red indicates *p* < 0.05; Dunnett’s *post-hoc* test, GFP as control. (**B, D**) Effect size and bootstrap 95% confidence interval of the data in A and C, respectively.

**Figure 5.**
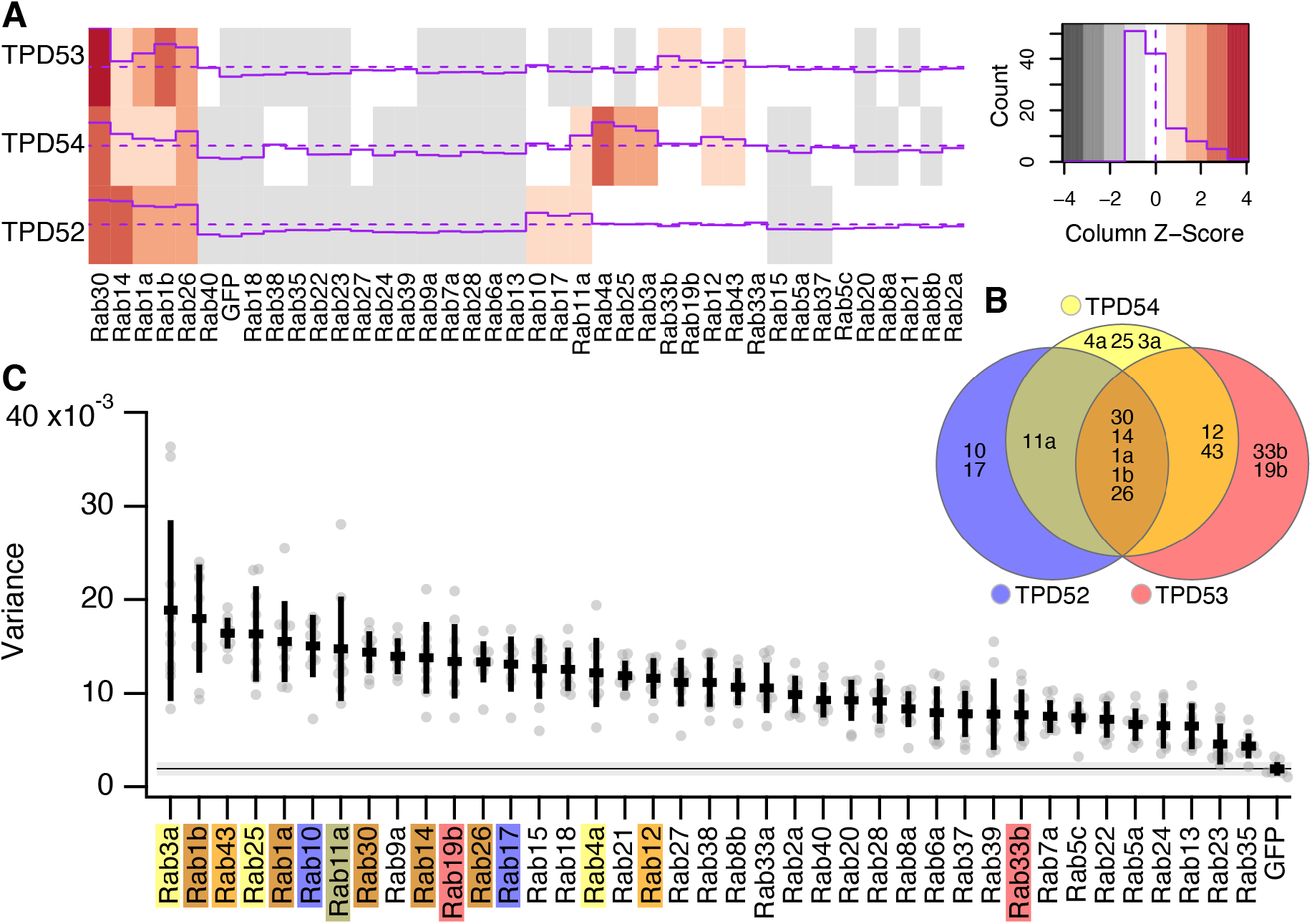
Spatiotemporal variance of GFP-Rab proteins. (**A**) Heat map showing the Rab screen data for TPD52, TPD53 (from Figure 4A,C), TPD54 (Larocque et al., 2020). Z-score of the values is color-coded and depicted as a purple line. Distribution of Z-values is shown in the color key. (**B**) Euler plot to show the Rabs identified in the screens and how they are linked to TPD52-like proteins. (**C**) Scatter dot plot of variance of fluorescence for the indicated GFP-Rab proteins expressed in HeLa cells. Rabs identified as present on INVs by hierarchical clustering of vesicle capture screens are indicated by the colours indicated in B. Dots represent individual cells, bars indicates the mean ±SD. The mean ±SD for GFP (control) is also shown as a black line and gray zone, across the plot.

The GFP-Rabs that were top hits in the vesicle capture screens all had subcellular distributions at steady state which were similar to that of TPD52-like proteins (Supplementary Video SV3). This suggested that if we quantify the spatiotemporal variance of individual Rabs, we could determine which Rabs are associated with INVs, to verify the results from the vesicle capture screens. Broadly, the Rabs that co-rerouted with one or more TPD52-like protein had high spatiotemporal variance and those that did not had lower variance (Figure 5C). This analysis verified that these Rabs were present on INVs that were positive for TPD52-like proteins. In theory this approach could also be used to identify INVs which lacked TPD52-like proteins. Indeed, high variance was seen for Rab18, Rab9a and Rab15 although none of these Rabs were co-rerouting with any of the three TPD52-like proteins tested (Figure 5C). Close inspection of these movies however suggest that the high variance of these Rabs is likely a false positive, since they are present on larger structures and not subresolution vesicles.

In summary this analysis suggests that there are four INV populations. The first population has all three TPD52-like proteins, and either Rab30, Rab14, Rab1a, Rab1b or Rab26. The second population is identified by a predominance of TPD52 and Rab10 or Rab17. The third, by the predominance of TPD54 and Rab4a, Rab25, or Rab3a. The fourth, by TPD53 and Rab33b or Rab19b. There are also intermediates: TPD52 and/or TPD54 with Rab11a and TPD53 and/or TPD54 with Rab12 or Rab43 (Figure 5B). Taken together, the data highlight the existence of different populations of INVs, marked by various Rabs, but all characterized by the presence of at least one member of the TPD52-like protein family.

### Amplification of TPD52-like proteins in cancers and potential changes in cell migration

Previous work showed that TPD52-like proteins are overexpressed in several cancer types (Nourse et al., 1998; Byrne et al., 1996), which may be associated with a more metastatic phenotype (Roslan et al., 2014; Shehata et al., 2008; Ummanni et al., 2008; Mukudai et al., 2013). To provide a full picture of TPD52-like protein overexpression in cancer, we analyzed the TCGA PanCancer Atlas (Supplementary Figure S3). Amplification of TPD52 and TPD54 was seen in a range of cancers including breast invasive carcinoma, ovarian and uterine cancers as well as cancers of the colon and liver. Amplification of TPD53 and TPD55 was less common (Supplementary Figure S3A). Rabs that were associated with INVs and those that were not were also analyzed. Of these, the INV-associated Rab25 had a similar amplification profile to TPD52-like proteins (Supplementary Figure S3B,C). Analysis of the ovarian serous carcinoma dataset showed that of 398 patients, amplification of TPD54 was seen in 29 (7%), amplification of Rab25 in 20 (5 %), and a significant co-occurrence in 7 patients (Log2 odds ratio >3, *p* < 0.001, *q* < 0.001. These results prompted us to investigate a potential link between TPD52-like proteins and cell migration and invasion.

### TPD52-like proteins are important for 2D and 3D cell migration

We imaged control RPE1 cells, and those depleted of TPD54, as they migrated on two different 2D substrates, fibronectin (Figure 6) or laminin (Figure S4). Tracking cells over 12 h, allowed us to generate a complete assessment of their migratory behavior (Figure S5). We found that TPD54-depleted cells had a strong reduction in migration speed on both substrates, indicating that this defect is not restricted to a single integrin heterodimer (Huttenlocher and Horwitz, 2011) (Figures 6A and S4). In addition, the directionality ratio over time changed to a slightly higher degree in TPD54-depleted cells compared with controls, suggesting a more erratic migratory behavior (Figure 6B).

**Figure 6.**
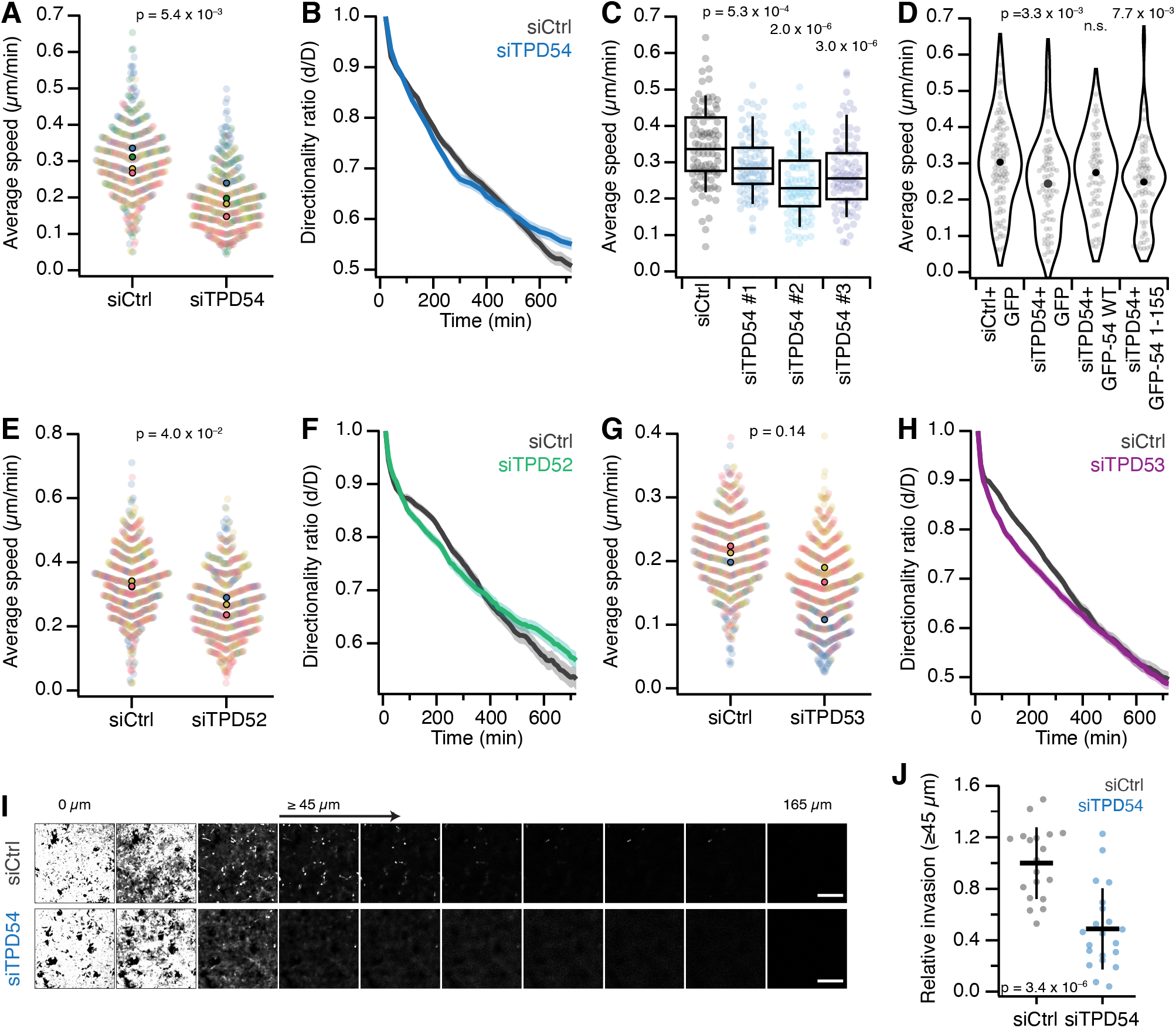
Role for TPD52-like proteins in cell migration and invasion. (**A-H**) RPE1 cell migration on a flat fibronectin substrate. (**A,E,G**) Superplots showing migration speed of control vs TPD54-(A), TPD52-(E) or TPD53-depleted (G) RPE1 cells migrating on a fibronectin substrate. Dots represent individual cells, color-coded for experiments. Markers, mean speed for individual experiments. P-value, Student’s T-test; *n*_exp_ = 4 (TPD54), 3 (TPD52 and TPD53); whereas *n*_cell_ = 469 (TPD54), 564,573 (TPD52), 588,597 (TPD53). (**B,F,H**) Graphs showing the directionality ratio over time of TPD54-depleted (B), TPD52-depleted (F) or TPD53-depleted cells (H). (**C**) Boxplots to show the migration speed of RPE1 cells on fibronectin and treated with siCtrl or each one of three TPD54-targeting siRNAs. Boxes show IQR, bar represents the median and whiskers show 9^th^ and 91^st^ percentiles. P-values from Dunnett’s *post-hoc* test using siCtrl as control; *n*_cell_ = 88 – 99, *n*_exp_ = 1. (**D**) Violin plot showing the average speed of siCtrl- or siTPD54-treated cells expressing GFP, GFP-TPD54 WT or GFP-TPD54 1-155, migrating on fibronectin. Dots, individual cells; Markers, mean speed. P-values from Dunnett’s *post-hoc* test using siCtrl + GFP as control. *n*_cell_ = 65 – 103, *n*_exp_ = 1. (**I-J**) Invasion of A2780 cells stably expressing Rab25 in a 3D context. Representative confocal images of siCtrl- or siTPD54-treated A2780 cells stably expressing Rab25 migrating through fibronectin-supplemented collagen type-I matrix for 48 h (I). Scale bar, 250 μm. Quantification of A2780 cell invasion in confocal sections ≥45μm, normalized to siCtrl (J). Dots, individual wells, Bars, mean ±SD. P-value, Student’s t-test.

To check whether the TPD54 RNAi phenotype was not the result of an off-target effect, we assessed migration speed on fibronectin of cells treated with a further two TPD54-targeting siRNAs (Figure 6C). A similar reduction was seen with all three siRNAs compared with control. We also tested whether the TPD54 RNAi phenotype could be rescued. We compared cells transfected with siCtrl expressing GFP with those transfected with siTPD54 that re-expressed GFP or GFP-TPD54 WT (Figure 6D). The migration defect was indeed rescued by re-expression of GFP-TPD54, but not GFP alone. More importantly, we wanted to test if the ability to bind to INVs was necessary for the role of TPD54 in migration. TPD54-depleted cells re-expressing a construct which lacked the INV-binding region (GFP-TPD54 1-155) failed to rescue the migration defect (Figure 6D). This shows that it is the ability to bind INVs that allows TPD54 to rescue the migration phenotype, and implicates INVs in cell migration.

Since we had established that TPD52 and TPD53 are also associated with INVs, we hypothesized that their depletion would also affect migration of cells on fibronectin (Figure 6E-H and Supplementary Figure S6). When compared with controls, TPD52-depleted RPE1 cells also showed a significant reduction in migration speed as well as an alteration in directionality ratio over time (Figure 6E-F). Similarly, TPD53-depleted cells show a decrease in speed and directionality, but there was more variation between experiments, which suggests a minor role for TPD53 in cell migration, in comparison with TPD52 and TPD54, which likely reflects their relative abundance (Figure 6G-H). In summary, TPD52-like proteins have a role in 2D cell migration and INVs are implicated in this function.

There are important differences between cells migrating on a flat substrate and those invading a 3D structure, and the latter is a more accurate model for cell movement in a cancer context. We therefore wanted to determine if TPD54 was also important for cellular invasion. A2780 ovarian carcinoma cells that stably express Rab25 are an accurate model of an aggressive tumor (Cheng et al., 2004), and Rab25 increases invasion in a 3D microenvironment. We depleted TPD54 using RNAi and allowed the cells to invade a fibronectin-enriched collagen matrix for 48 h (Figure 6I). We quantified the invasion by measuring the area that cells occupy above 45 μm, and found that TPD54-depleted cells lose their ability to invade a dense 3D matrix (Figure 6J). Taken together, the data show that TPD54 is important for cell migration, both in non-cancerous RPE1 cells migrating on a 2D substrate, and in cancer cells invading a 3D structure.

### INVs are involved in integrin trafficking

In addition to changes in cell motility, we observed that TPD54-depleted cells seemed morphologically different from control cells (Figure 7A). To quantify this difference, we analyzed a series of cell shape parameters using a semi-automated workflow (see Methods, Figure S7). TPD54-depleted cells had much larger footprints, with an average area that was almost twice that of control cells (Figure 7B). Close inspection of movies of cells migrating on fibronectin revealed that the larger area and reduced speed of migration were linked. TPD54-depleted cells behaved as if they were “stuck” to the substrate; instead of having one clear lamellipodium, they made several smaller ones (Supplementary Video SV4). These observations are consistent with changes in directionality as seen in Figure 6B. We hypothesized that these results were the result of a defect in integrin trafficking. Therefore, we assessed the ability of TPD54-depleted RPE1 cells to recycle integrins. Using an integrin *α*5 recycling assay, we found that TPD54-depleted cells show a marked reduction in recycling of integrins compared to control cells (Figure 7C). These data confirm that TPD54 and the INVs are involved in integrin traffic and provide a mechanistic explanation for the cell migration and invasion phenotypes observed in cells depleted of TPD52-like proteins.

**Figure 7.**
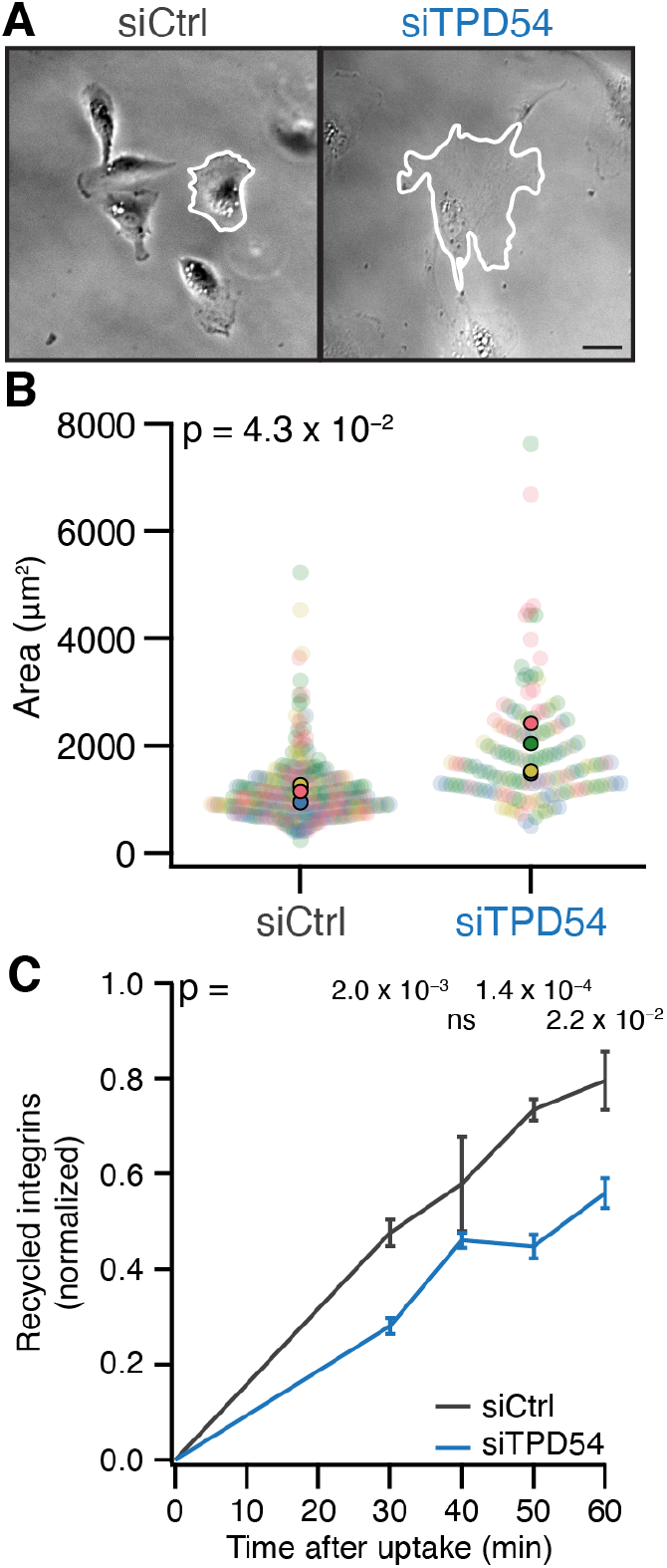
TPD54-depleted cells show defective integrin trafficking. (**A**) Example micrograph of control and TPD54-depleted RPE1 cells migrating on fibronectin. The perimeter of the cell is outlined (white). Scale bar, 10μm. (**B**) Violin plot showing the area of control vs TPD54-depleted RPE1 cells migrating on fibronectin. Dots represent individual cells, markers show the median. p value from Student’s test. *n*_cell_ = 288 – 151, *n*_exp_ = 4. (**C**) Quantification of integrin *α*5 recycling over time in siCtrl (gray line) or siTPD54-treated (blue line) RPE1 cells. Bars show SD. *n*_exp_ = 3.

## Discussion

In this study, we demonstrated the involvement of INVs in integrin trafficking and cell migration. We described how the TPD52-like protein TPD54 is localized to INVs and showed that other members of this family, TPD52 and TPD53, are also INV proteins. This allowed us to document the Rab GTPase complement of INVs, which included Rabs that are involved in integrin traffic and cell migration. Depletion of TPD52-like protein family members caused decreased cell migration and invasion, a phenotype that could be linked to decreased integrin recycling.

The binding of TPD52-like proteins to INVs can be explained by two molecular properties. First, preference for high curvature membranes. INVs are ~30nm in diameter (Larocque et al., 2020) and we found that TPD54 binds 30 nm or 50 nm liposomes more efficiently than those of 100 nm or 200 nm diameter. This reflects the subcellular distribution of TPD52-like proteins in cells, which are found predominantly on INVs but also on endosomes and Golgi. Second, binding is via positively charged residues in a region (residues 155-180, human TPD54 numbering) that is conserved across metazoans. A single charge-swap mutation in this region was sufficient to reduce liposome binding or association with INVs in cells. General ionic interactions between the TPD52-like protein and the negatively-charged bilayer rather than a phosphoinositide preference, may also explain how these proteins can bind INVs, which are likely to have a variety of lipid compositions (van Meer and de Kroon, 2011). It was surprising that the curvature preference *in vitro* was also lost by the charge-swap mutation suggesting that these two molecular properties are linked. This recipe of molecular properties is reminiscent of BAR domain-containing proteins such as amphiphysin that are crescent-shaped dimers with positive charges on the inner surface (Peter et al., 2004). However, we found that disruption of TPD54 multimerization did not affect its ability to bind INVs, which suggests that these proteins may have a different mode of binding. Answering precisely how TPD52-like proteins bind INVs will require a high resolution structure of a TPD52-like protein.

We exploited the direct binding of TPD52-like proteins to INVs in a mitochondrial vesicle capture assay in order to detail the collective Rab GTPase complement of INVs. This analysis showed that the TPD52-like proteins associate with similar INVs as delineated by their Rab GTPases. The Rabs associated with INVs cover anterograde, Golgi and recycling trafficking, with the addition of Rab21, an endocytic Rab, which was captured using TPD52 only (Simpson, 2004). This Rab is notable here since it is involved in the internalization of integrins (Pellinen et al., 2006). Rab30, a Golgi resident Rab that is not fully characterized, was our top hit in vesicle capture using TPD52, TPD53 or TPD54; by this definition it could be considered as an “INV Rab”. Among the Rab30 effectors identified in a screen in *Drosophila*, were the dynein adaptor BicD, and the tethering factors GARP and exocyst (Gillingham et al., 2014), suggesting that Rab30 is involved in membrane fusion between the endosomes and the Golgi, and between the Golgi and the plasma membrane. Rab30 has been shown to bind to PI4KB in an autophagy context (Oda et al., 2016; Nakajima et al., 2019). Whether Rab30 and PI4KB are needed for the formation or transport of INVs, or whether GARP and exocyst are tethering factors present on INVs remain to be determined. Although limited to the 39 Rabs tested, we found no evidence for Rab-positive INVs that had no TPD52-like protein. This suggests that TPD52-like proteins are core components of INVs and possibly should be considered the molecules that define this class of transport vesicle.

We found that depletion of TPD54, TPD52, and, to a lesser extent TPD53, decreased cell migration. This mirrored the literature showing that TPD52-like proteins proteins are overexpressed in various cancers and that overexpression is potentially correlated with a more invasive, migratory phenotype (Roslan et al., 2014; Shehata et al., 2008). INVs are implicated in cell migration not only because depletion of a core INV protein impaired migration speed but also because normal motility could only be rescued by wild-type TPD54 and not by a truncated form that cannot bind to INVs. Of the Rabs that we found associated with INVs, Rab4, Rab11 and Rab25 are all involved in integrin trafficking and cell migration (Caswell et al., 2007; Powelka et al., 2004; Roberts et al., 2001). After internalization, the integrin heterodimers are recycled back to the plasma membrane via a short, Rab4-dependent pathway or a long, Rab11-dependent pathway (Roberts et al., 2001); and in cancer cells, Rab25 sorts ligand-free integrins for recycling at the leading edge and ligand-bound integrins to lysosomes where they reach the plasma membrane and cell rear in a CLIC3-dependent manner (Dozynkiewicz et al., 2012). Previously, we found that recycling of internalized transferrin was affected by depletion of TPD54, implicating INVs in receptor recycling (Larocque et al., 2020). Here, we showed that recycling of *α*5 integrin is impaired when TPD54 was depleted, suggesting that INVs also mediate recycling of internalized integrins. This raises an interesting question: if the size of INVs in invariant (~30nm diameter), how many integrin heterodimer can be trafficked in an INV? When the integrins are recycled to the plasma membrane, they are in their bent, inactive conformation. For integrin *α*IIb*β*3, this conformation extends 11 nm (Ye et al., 2008), which indicates that traffic in INVs is possible. Despite their small size, the maximum capacity of an INV is surprisingly high (Martins Ratamero and Royle, 2019), although for bulky cargoes such as integrins, the number traveling in each INV is likely to be low.

## Methods

### Molecular biology

Several plasmids were available from previous work, e.g. to express mCherry-FKBP-TPD54, GFP-FKBP-TPD54, mCherry-MitoTrap, dark MitoTrap (pMito-dCherry-FRB) (Larocque et al., 2020). All TPD52-like protein constructs used in the paper represent the longest isoform (splice isoform 1). To make mCherry-FKBP-TPD52, human TPD52 (synthesized by GeneArt) was inserted in place of TPD54 in mCherry-FKBP-TPD54 using BglII and MfeI. FLAG-TPD52 was made by amplifying human TPD52 from the synthesized gene and inserting into pFLAG-C1 via BglII and MfeI. To make mCherry-FKBP-TPD53, TPD53 (GeneArt synthesis) was inserted into pmCherry-FKBP-C1 via HindIII and BamHI. FLAG-TPD53 was made by amplifying TPD53 from the synthesized gene and inserting into pFLAG-C1 via BglII and BamHI. The plasmid to co-express untagged TPD54 and GFP (pIRES-EGFP-TPD54) was made by amplifying TPD54 by PCR from human Tumor protein D54 (IMAGE clone: 3446037) and inserting into pIRES-EGFP-puro (Addgene #45567) via NheI and XhoI. The mCherry-FKBP-TPD54 deletions (1-37, 1-82, 38-82, 83-206, 1-155, 1-1801-37, 1-82, 38-82, 83-206, 1-155, 1-180) were made by PCR from mCherry-FKBP-TPD54 and were each inserted into pmCherry-FKBP-C1 via BglII and MfeI. The mCherry-FKBP-TPD54 mutants (K15E, R95E, K154E, R159E, K165E and K175E, K177E) and GFP-FKBP-TPD54 mutant (L53P, L67P) were created by site-directed mutagenesis (SDM). Plasmids to express GFP-tagged Rabs were a gift from Francis Barr (University of Oxford, UK), except for GFP-Rab1a and GFP-Rab5c, which were described in (Larocque et al., 2020).

### Cell culture

HeLa cells (Health Protection Agency/European Collection of Authenticated Cell Cultures, #93021013) were maintained in DMEM supplemented with 10 % FBS and 100 U ml^-1^ penicillin/streptomycin. RPE1 cells (HD-PAR-541 clone 7724) were maintained in Ham’s F12 Nutrient Mixture supplemented with 100 U ml^-1^ penicillin/ streptomycin, 10% FBS, 2.3g/l sodium bicarbonate and 2 mM L-Glutamine. A2780 human ovarian cancer cells (female) stably expressing Rab25 (Caswell et al., 2007) were maintained in RPMI-1640 medium (Sigma) supplemented with 10 % FCS, 1 % L-Glutamine and 1 % antibiotic-antimycotic (Sigma). All cell lines were kept at 37 °C and 5% CO_2_. RNA interference was done by transfecting 100 nM siRNA (TPD54#1: 5’-GUCCUACCUGUUACGCAAU-3’, TPD54#2: CUCACGUUUGUAGAUGAAA, TPD54#3: CAUGUUAGCCCAUCAGAAU; TPD52: 5’-CAAAUAGUUUGUGGGUUAA-3’; TPD53: 5’-GUCUCCAGCAAUAGGAUGAUUUACUA-3’) with Lipofectamine 2000 (Thermo Fisher Scientific) according to the manufacturer’s protocol. For DNA plasmids, cells were transfected with a total of 600 ng DNA (per well of a 4-well LabTek dish) using 0.75 μl Genejuice (Merck Millipore) following the manufacturer’s protocol. The A2780 cells were transfected by electroporation using a nucleofector (Amaxa, Lonza) using solution T, program A-23, 20 nM siRNA as per the manufacturer’s instructions. Invasion experiments were performed 24h post nucleofection.

### Biochemistry

For immunoprecipitation, HeLa cells were seeded in 10 cm dishes. Either FLAG-TPD54, FLAG-TPD53 or FLAG-TPD52 was transfected with either GFP-FKBP, GFP-FKBP-TPD54 or GFP-FKBP-TPD54 L53P, L67P, 10 μg total using GeneJuice (Merck Millipore) according to the manufacturer’s protocol. After 48 h, the cells were lysed with 10 mM pH 7.5 Tris, 150 mM NaCl, 0.5 mM EDTA, 1 % Triton X-100 and protease inhibitors. Lysate was passed through a 23G syringe and spun in a benchtop centrifuge for 15 min at 4 °C. The cleared lysate was then incubated for 2h at 4 °C with 10 μl of anti-FLAG M2 Magnetic beads (Sigma-Aldrich, M8823) pre-washed with TBS. The beads were then washed 3 times with cold TBS, resuspended in Laemmli buffer and run on a precast 4 to 15 % polyacrylamide gel (Bio-Rad).

For western blotting, the following primary antibodies were used: rabbit anti-TPD54 (Dundee Cell products), 1:1000; goat anti-TPD53 (Thermo, #PA5-18798), 0.5μg/ml; mouse anti-GFP clones 7.1 and13.1 (Sigma Roche: 118144600010), 1:1000; and mouse anti-FLAG M2 (Sigma-Aldrich, F1804), 1 μg/ml. HRP-conjugated secondary antibodies were used with enhanced chemiluminescence detection reagent (GE Healthcare) for detection and manual exposure of Hyperfilm (GE Healthcare) was performed.

### Protein expression and purification

GST, GST-TPD54 and GST-TPD54 R159E were expressed in *E. coli* BL21 cells grown in double yeast tryptone media. Starter cultures of 10 ml were grown overnight at 37°C and shaken at 200 rpm. They were then diluted into 400 ml cultures and grown at 37°C, 200 rpm until an optical density at 600 nm between 0.6 and 0.8. To induce expression of the proteins, IPTG was added to a final concentration of 0.5 mM and cells were grown for a further 5 h at 37°C, 200 rpm. The cells were harvested by centrifugation at 9200 *g* for 10 min at 4 °C, washed with cold PBS and pelleted again at 3200 *g* for 15 min at 4 °C. Pellets were stored at −80 °C until purification.

For purification, pellets were resuspended in 50 ml lysis buffer (50 mM Tris, 150 mM NaCl, protease inhibitor cocktail tablet (Roche), 0.2 mM PMSF, pH 8) and lysed by sonication. Cell debris was pelleted by centrifugation at 34600 *g* for 30 min at 4°C. The supernatant was loaded onto a GSTrap column (GE Healthcare), which was then washed with 10 column volumes of wash buffer (50 mM Tris, 150 mM NaCl, protease inhibitor cocktail tablet, 0.1 mM PMSF, pH 8), then 10 column volumes of high salt wash buffer (50 mM Tris, 500 mM NaCl, protease inhibitor cocktail tablet, 0.1 mM PMSF, pH 8). The GST-tagged proteins were eluted by the addition of elution buffer (50 mM Tris, 150 mM NaCl, 50 mM glutathione, pH 8). Purified proteins were dialysed into binding buffer (150 mM NaCl, 20 mM HEPES, pH 7) for use in liposome binding assays.

### Protein-Liposome interactions

Folch extract from bovine brain (Sigma-Aldrich) was dissolved in chloroform. The lipid mixture was dried with nitrogen flow followed by 2h desiccation in a vacuum. The dried lipids were resuspended in binding buffer (150 mM NaCl, 20 mM HEPES, pH 7) to a final concentration of 4mM. To generate liposomes with desired diameters, the lipid mixture was heated to 60°C and extruded 15× through polycarbonate membrane filters (Avanti Polar Lipids) with pore sizes of 200, 100, 50 or 30 nm. For 30 nm and 50 nm sizes, liposomes were produced by first extruding through a 100 nm filter, followed by extrusion through filters of a smaller pore size. Note that material is lost with each extrusion. Liposomes were stored at 4 °C until used in binding assays.

For the liposome pelleting assay, liposomes were centrifuged at 100 000 *g* for 25 min at 4 °C. Pellet and supernatant samples were prepared with NuPAGE LDS Sample Buffer + reducing agent (Invitrogen) and incubated at 70°C for 10 min. Lipids were resolved on 12 % Bis-Tris gels run in MES buffer and stained with 0.1% Coomassie in 10% acetic acid for 5 min (method adapted from Boucrot et al. (2012)). Gels were destained in water overnight to leach the loading dye.

For the protein-liposome binding assay, purified GST-tagged proteins were pre-cleared before use and then protein (2 μM) and liposomes (600 μM) were mixed with binding buffer (150mM NaCl, 20mM HEPES, pH 7) to a total volume of 200 μl and incubated on ice for 20 min. Samples were then centrifuged at 100 000 *g* for 25 min at 4 °C. Bound (pellet) and free (supernatant) samples were prepared for SDS-PAGE by adding Laemmli buffer and boiling for 5 min. Proteins were resolved by SDS-PAGE on 4% to 15% Mini-PROTEAN TGX gels and visualized by staining with InstantBlue (Expedeon).

### Integrin recycling assay

The ELISA plate (maxisorp 96 wells, Thermo Scientific) was prepared the day before the experiment by incubating the wells with 50μl/well of 5 μg/ml of anti-integrin *α*5 antibodies (BD Biosciences, 555651) in 0.05m Na_2_CO_3_ pH 9.6, overnight at 4°C. 10cm dishes were seeded with RPE1 cells, in triplicate. Cells were serum starved for 30 min at 37°C. Following two 5 ml washes with cold PBS, the surface receptors were labeled with 0.133 mg/ml of EZ-Link Sulfo-NHS-SS-Biotin (Thermo scientific, 21331) in PBS at 4 °C for 30 min. Cells were washed twice with 5 ml of cold PBS on ice and 5 ml of warm serum-free medium was added. Plates were incubated at 37 °C for 30 min to allow receptor internalization and then washed again with 5 ml of cold PBS on ice.

The cells were washed with cold reduction buffer (50 mM Tris pH 7.5, 102.5 mM NaCl, pH 8.6). Cell surface was reduced by adding 3 ml reduction buffer and 1 ml of Mesna buffer (390 mg of Mesna was added to 26 ml of reduction buffer, mixed thoroughly and 39μl of 10m NaOH was added). Plates were agitated at 4 °C for 20 min, then washed twice with cold PBS. The plates were then incubated in warm medium at 37°C to allow receptor recycling. The cell surface was then reduced again for 20 min as described above. Reduction buffer containing iodoacetamide (442 mg in 26 ml PBS) was added to the reduction buffer (1:4) to quench the reaction, for 10 min. The ELISA plate was blocked with 5% BSA in PBS-Tween at room temperature for 1 h. The cells were washed twice with cold PBS on ice. Lysates were obtained by scraping the cells with a total of 100 μl/condition of lysis buffer (200 mM NaCl, 75 mM Tris pH 7.5, 15 mM NaF, 1.5 mM Na_3_VO_4_, 7.5 mM EDTA, 7.5 mM EGTA, 1.5% Triton-X100, 0.75% Igepal, protease inhibitors). The ELISA plate was washed twice with PBS-Tween and 50 μl of lysate was put in each well, covered with parafilm and incubated overnght at 4 °C. Following 5× washes with PBS-Tween, 50 μl of 1 μg/ml streptavidin-HRP, 1 % BSA in PBS-Tween was added to each well and incubated for 1 h at 4 °C. Five more washes were performed with PBS-Tween and 50 μl of detection reagent (0.56 mg/ml of ortho-phenylenediamine dihydrochloride in ELISA buffer (25.4 mM NaHPO_4_, 12.3 mM citric acid, pH 5.4) with 0.003% H_2_O_2_) was added to each well. The plate was incubated at room temperature in the dark for 15 min, before reading on a plate reader using 450 nm light.

### Microscopy

For confocal imaging, cells were grown in 4-well, glass-bottom, 3.5cm dishes (Greiner Bio-One), and medium was exchanged for Leibovitz L-15 CO_2_-independent medium for imaging at 37°C on a spinning disc confocal system (Ultraview Vox, PerkinElmer) with a 100 × 1.4 NA oil-immersion objective. Images were captured using an ORCA-R2 digital charge-coupled device camera (Hamamatsu) following excitation with 488 nm and 561 nm lasers. Rerouting of mCherry-FKBP-TPD52 or mCherry-FKBP-TPD53 to the mitochondria (dark MitoTrap) was induced by addition of 200 nM rapamycin (Alfa Aesar). For the Rab GTPase co-rerouting experiments, an image before rapamycin and an image 2min after rapamycin were taken of live cells. For the flicker analysis, cells were imaged for 30 frames with a 300 ms exposure.

For the invasion assay, 5 mg/ml collagen I supplemented with 25 μg/ml fibronectin was polymerized in inserts (Transwell; Corning) at 37°C for 1 h. The inserts were inverted and A2780 cells stably expressing Rab25 were seeded on the opposite side of the filter. The inserts were then put in 0.1 % serum medium supplemented with 10 % FCS and 30 ng/ml EGF was added on top of the matrix. After 48 h, cells were stained with Calcein-AM and visualized with an inverted confocal microscope (TCS SP5 AOBS; Leica) using a 20× objective. Cells were considered invasive above 45 μm and slices were taken every 15 μm.

For widefield imaging, 4-well LabTek dishes were incubated 30 min at 37°C with 10 μg/ml fibronectin or O/N at 37°C with 20 μg/ml laminin (Sigma, L2020-1MG). RPE1 cells were then plated at low density in the dishes and imaged the next day, in L15 medium supplemented with 10 % FBS and 100 U ml^-1^ penicillin/streptomycin. Cells were imaged at 37°C on a Nikon Ti epifuorescence microscope, with a 20× 0.50NA air objective, a heated chamber (OKOlab) and CoolSnap MYO camera (Photometrics) using NIS elements AR software. Movies were recorded over 12 h (1 frame/10 min) with phase contrast.

### Correlative light-electron microscopy

Analysis of vesicle capture and mitochondrial aggregation was by correlative light-electron microscopy (CLEM) following the methods outlined previously (Larocque et al., 2020). Briefly, transfected cells were plated onto gridded dishes (P35G-1.5-14-CGRD, MatTek Corporation, Ashland, MA, USA). Cells were imaged at 37°C in Leibovitz L-15 CO_2_-independent medium supplemented with 10 % FBS. Rapamycin (200 nM, final concentration) was added for 3 min before the cells were fixed in 3% glutaraldehyde, 0.5% paraformaldehyde in 0.05M phosphate buffer pH 7.4 for 1 h. Aldehydes were quenched in 50 mM glycine solution, then post-fixed in 1% osmium tetroxide and 1.5% potassium ferrocyanide for 1 h and then in 1% tannic acid for 45 min to enhance membrane contrast. Cells were rinsed in 1% sodium sulphate then twice in H_2_O before being dehydrated in grade series ethanol and embedded in EPON resin (TAAB). Correlation of light images allowed the cell of interest to be identified for sectioning and 70 nm ultrathin sections were cut and collected on formvar-coated hexagonal 100 mesh grids (EM resolutions). Sections were post-stained with Reynolds lead citrate for 5 min. Electron micrographs were recorded using a JEOL 1400 TEM operating at 100 kV using iTEM software.

### Data analysis

To quantify liposome binding, the mean pixel density of gel bands was measured in Fiji. The background for each lane was subtracted and the precipitation (no vesicle control band) was subtracted from the bound sample bands. The bound band intensities (multiplied by 2, since 1/2 was loaded) was plotted as a percentage of the bound intensity plus the free band intensity (multiplied by 34, since 1/34 was loaded).

To quantify liposome pelleting, gel bands densitometry was as above. The amount of lipid pelleted for each liposome size was calculated as a percentage of 100 nm pelleting. The protein binding results for each liposome size were then normalized according to the amount pelleted to enable a fair comparison of the results across vesicle sizes.

For the Rab screen, co-rerouting of Rab GTPases was quantified by averaging for each cell, the pixel intensity in the green channel in 10 ROIs of 10×10 pixels on the mitochondria, before and after rapamycin. This mitochondrial intensity ratio (*F_post_/F_pre_*) for every Rab was compared to the ratio of GFP in TPD52- or TPD53-rerouted cells. Estimation statistics were used to generate the difference plot shown in Figure 4. The mean difference is shown together with bias-corrected and accelerated (BCa) 95% confidence intervals calculated in `R` using 1 × 10^5^ bootstrap replications. The heat map was generated in `R` (see Code Availability) using mitochondrial intensity ratio data. Note that, when constructing the heat map we reanalyzed data for Rab3a and Rab4a from the original TPD54 screen (Larocque et al., 2020) and this has slightly altered the Rab profile for TPD54.

Analysis of fluorescence variance in the GFP-Rab signal in live cell movies was done using 5 20 × 20 pixel excerpts of 30 frames from live cell imaging captured at 0.1775 s per frame. The excerpts were positioned in the cytoplasm to avoid measuring variation in endosomal, ER or Golgi structures. Each frame was first normalized to the mean pixel intensity for that frame, then the variance per pixel over time was calculated, resulting in a 20 × 20 matrix of variances. The mean of the 5 matrices is presented as the “variance” for that cell. The analysis of the fluorescence variance in the mCherry-FKBP-TPD54 mutants was measured in the same way, except that 1 ROI of 50 × 50 pixel was taken per cell.

Invasion was quantified using the area calculator plugin in Fiji, measuring the fluorescence intensity of cells invading 45 μm or more and expressing this as a percentage of the fluorescence intensity of all cells within the plug. The data was normalized to siCtrl, to show relative invasion.

For the 2D migration assay, cells were tracked using the Fiji plugin Manual Tracking by using the center of the nucleus as guide. The data were saved as csv files and fed into `CellMigration 1.13` in IgorPro for analysis (Royle, 2019).

For the cell shape analysis, a scientist blind to the experimental conditions drew with a stylus the outline of each cell in a frame halfway through the migration movies. The coordinates of all cell contours were fed into `CellShape 1.01` in IgorPro for analysis (Royle, 2020).

Figures were made with Fiji or Igor Pro 8 (WaveMetrics), and assembled using Adobe Illustrator. Null hypothesis statistical tests were as described in the figure legends.

### Data and software availability

All code that is specific to this manuscript is available at https://github.com/quantixed/p054p031.

## ACKNOWLEDGEMENTS

The authors thank Laura Cooper, Erick Martins Ratamero and Claire Mitchell of CAMDU (Computing and Advanced Microscopy Unit) for their support and assistance in this work. We would like to thank Francis Barr for reagents and Darius Koester for advice and assistance with liposome production. We would also like to thank Miguel Hernández González and Joseph Cockburn for valuable discussion. GL was supported by Fonds de recherche du Québec - Nature et technologies and University of Warwick Chancellor’s Award. The work was supported by the UK Medical Research Council (MR/P018947/1).

## AUTHOR CONTRIBUTIONS

G. Larocque performed experimental work, analyzed data, and wrote the paper. P.J. La-Borde performed the liposome binding assay. B.J. Wilson and P.T. Caswell performed the invasion assay. N.I. Clarke did the vesicle capture CLEM experiments. D.J. Moore helped with the Rab screens. S.J. Royle analyzed data, wrote computer code, and wrote the paper.

## COMPETING FINANCIAL INTERESTS

The authors declare no conflict of interest.

## Supplementary Information

**Figure S1.**
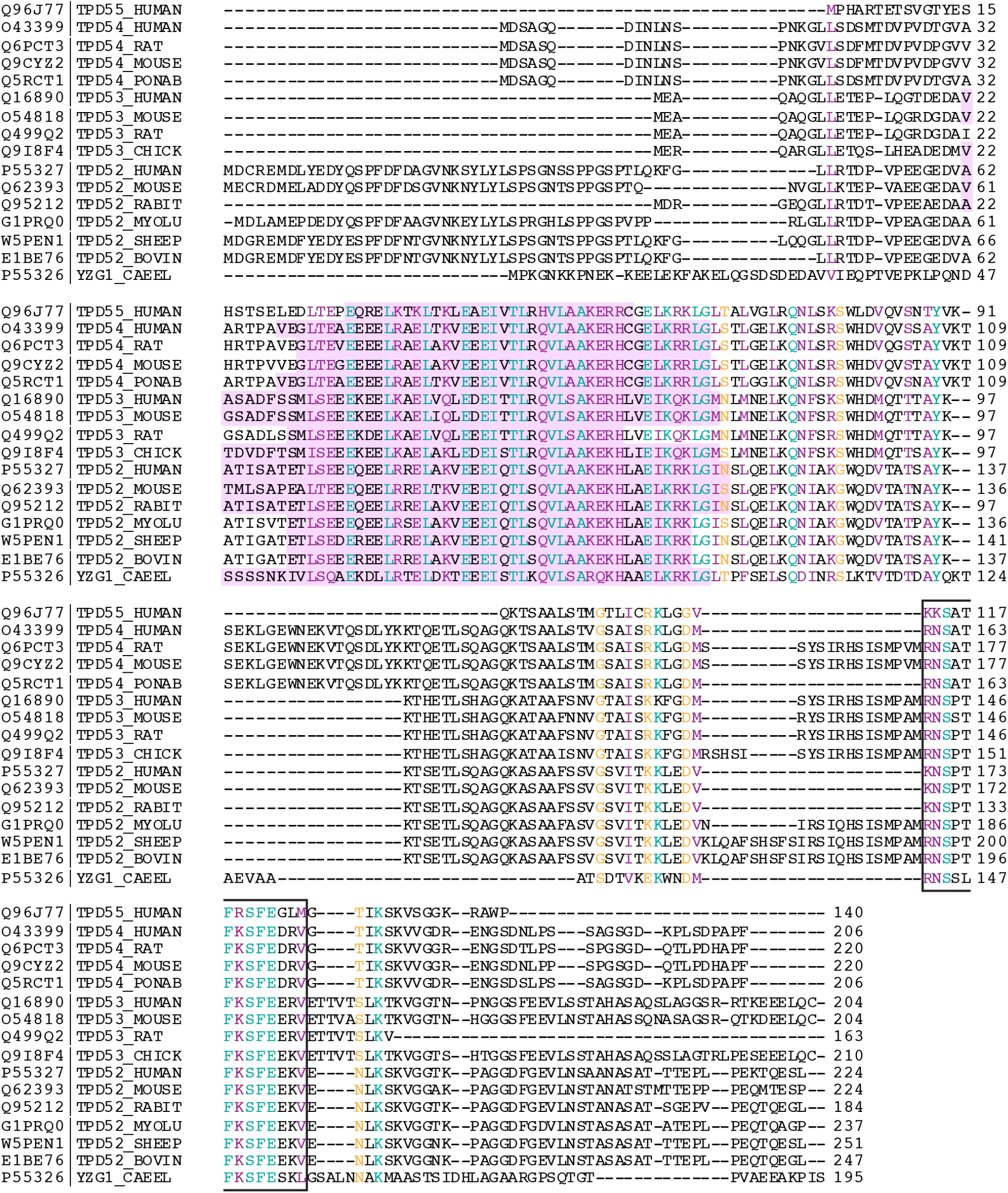
Alignment of the amino acid sequences of the TPD52 family members from different species. Alignment of TPD52-like protein family members from human, mouse, rat, orangutan, chicken, rabbit, bat, sheep, bovine and *C. elegans*. Pink highlighted area, coiled-coil domain. Boxed area, conserved region. Lettering: Teal, fully conserved residues; Purple, strongly similar (>0.5 in the Gonnet PAM 250 matrix); Yellow, weakly similar (<0.5 in the Gonnet PAM 250 matrix).

**Figure S2.**
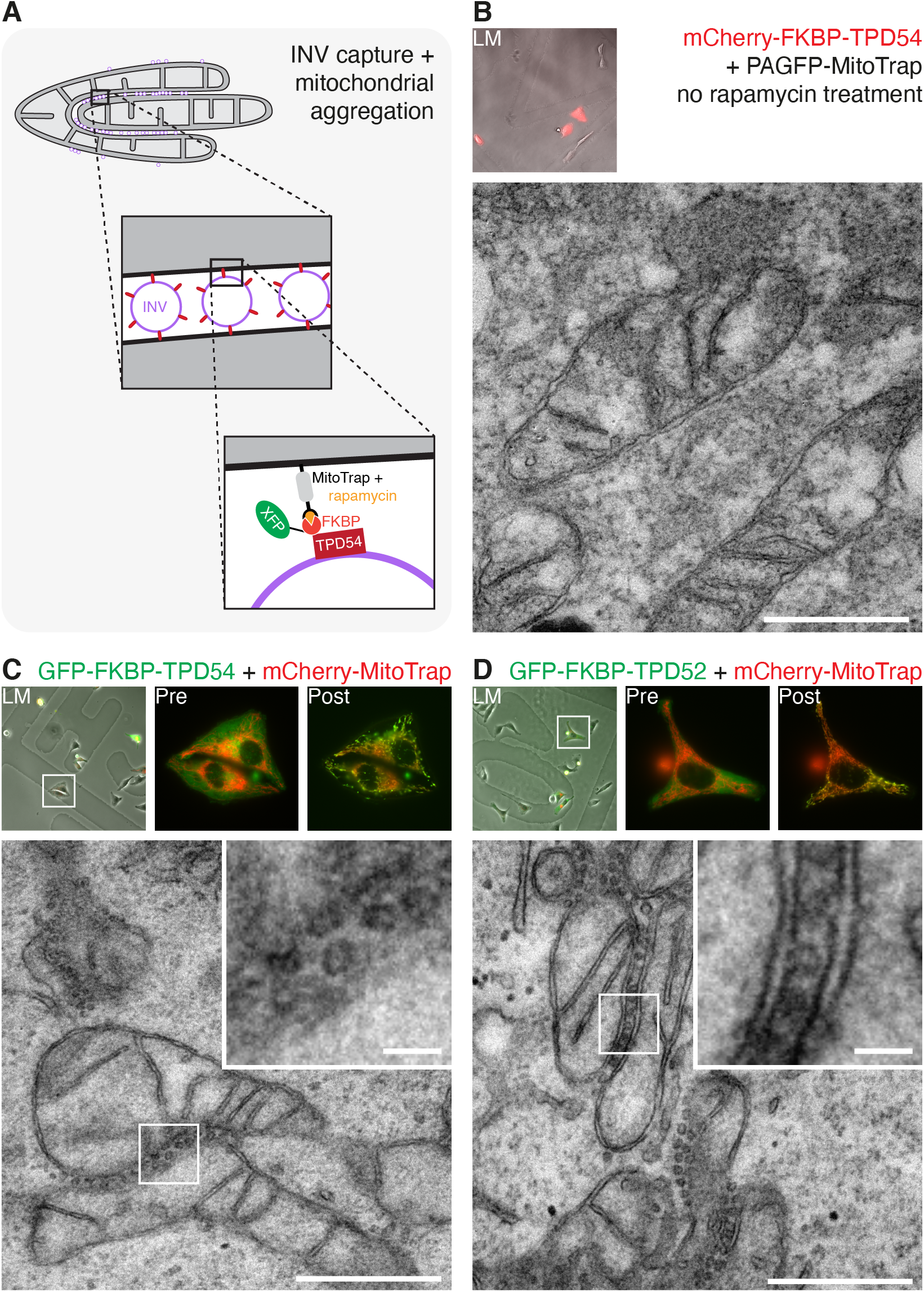
Vesicle capture and mitochondrial aggregation using TPD52 or TPD54. (**A**) Schematic diagram of vesicle capture at mitochondria and their subsequent aggregation. MitoTrap is an FRB domain targeted to mitochondria, FKBP-tagged TPD52 or TPD54 are co-expressed and, when rapamycin is added, the INVs associated with TPD52-like proteins become trapped at the mitochondria, eventually causing aggregation of mitochondria. (**B-D**) Correlative light electron microscopy (CLEM) experiments to test for vesicle capture. (**B**) Control. Cells expressing mCherry-FKBP-TPD54 and PAGFP-MitoTrap with no addition of rapamycin. No mitochondrial aggregation observed. (**C**) Cells expressing GFP-FKBP-TPD54 and mCherry-MitoTrap with rapamycin 200 nM for 3 min. The same cell is shown in Figure 2. (**D**) Cells expressing GFP-FKBP-TPD52 and mCherry-MitoTrap with rapamycin 200nM for 3min. Insets: 4× zoom. Scale bars, 500nm; insets, 50 nm.

**Figure S3.**
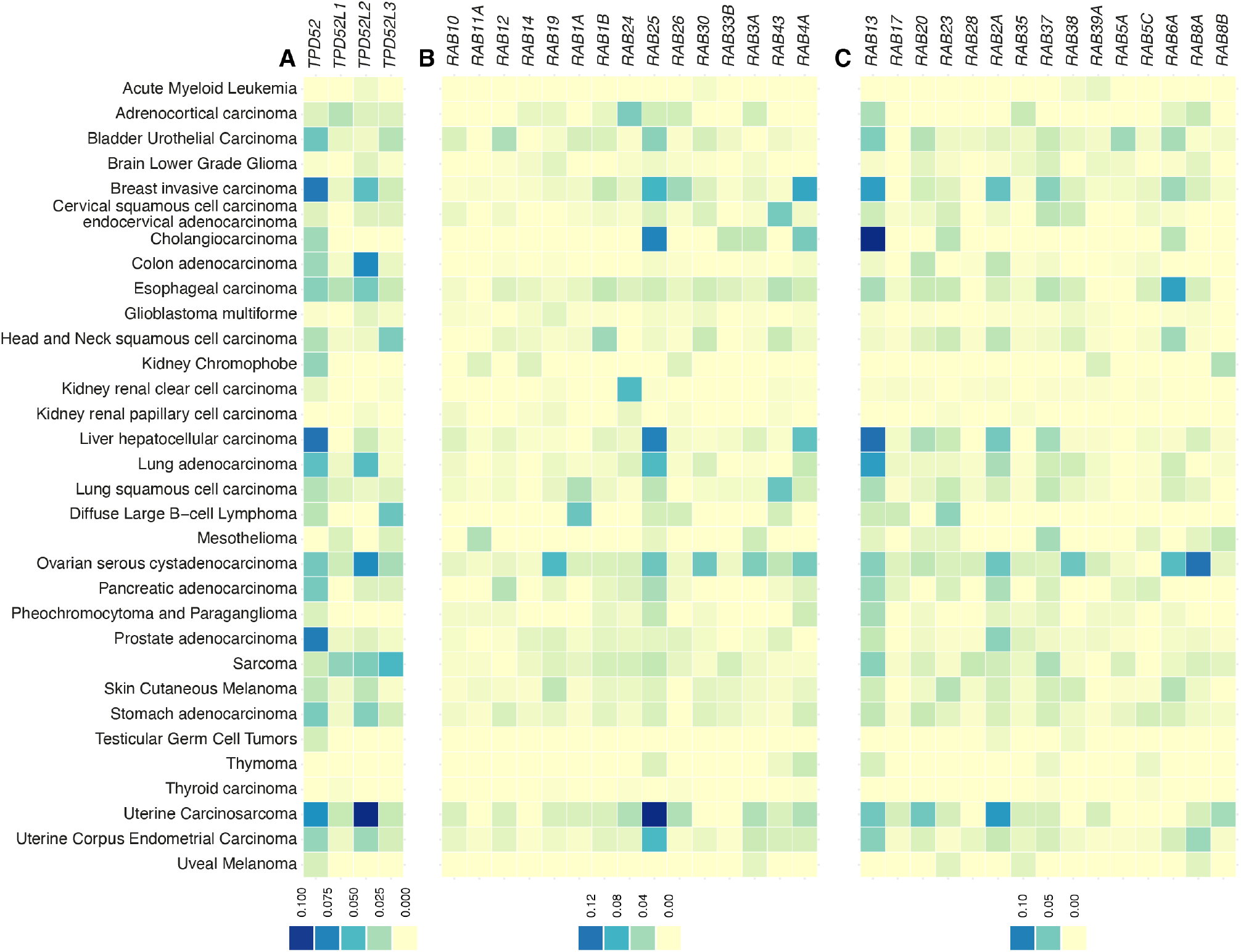
Amplification of TPD52-like proteins in cancer. Frequency of amplification of the indicated genes in TCGA PanCancer Atlas Studies (10967 samples). (**A**) TPD52-like proteins. (**B**) Rab GTPases associated with INVs. (**C**) Rab GTPases not associated with INVs. Colorscale indicates the fraction of samples of each cancer type showing amplification.

**Figure S4.**
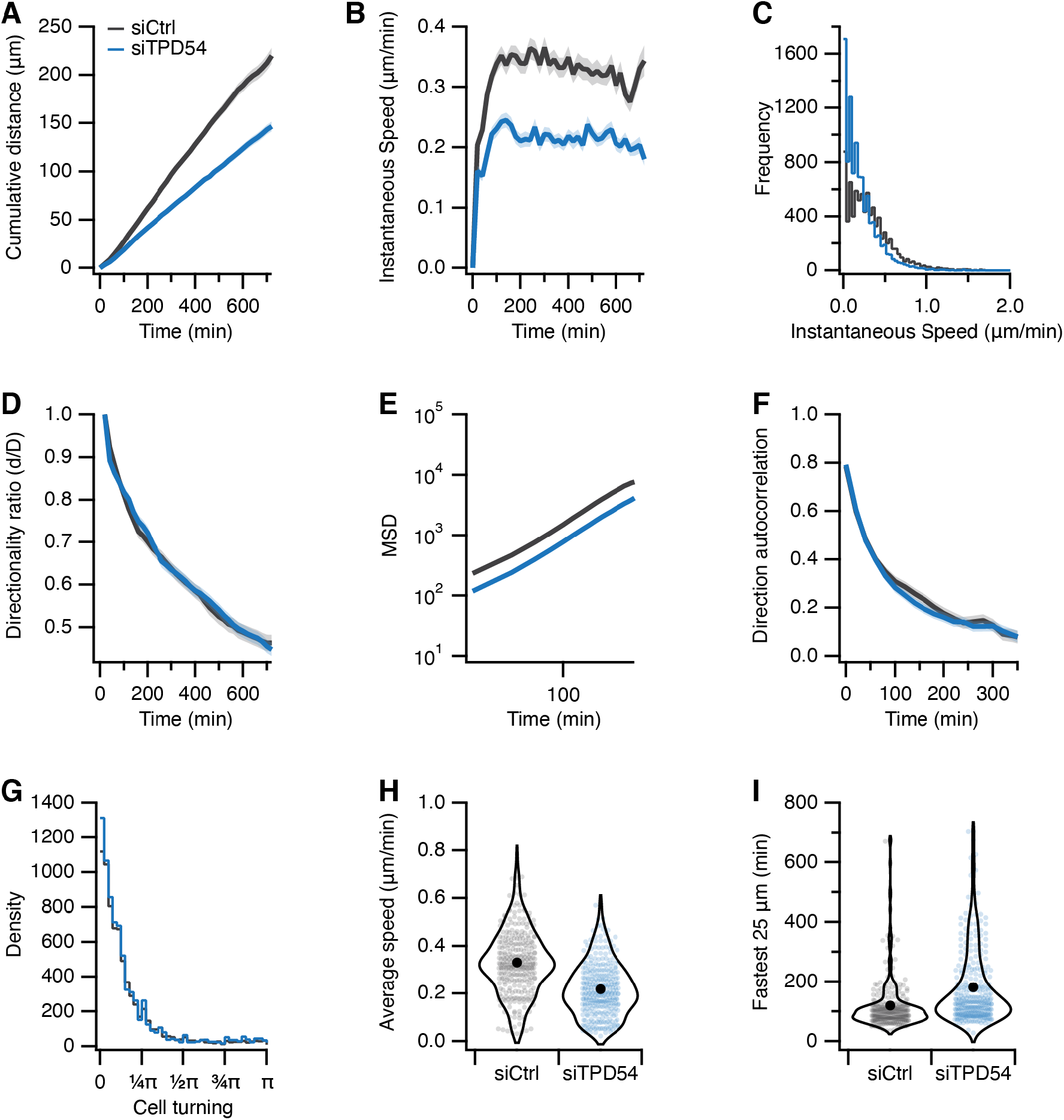
Migration of RPE1 cells on laminin. Contrl (gray) or TPD54-depleted (blue) RPE1 cells migrating on laminin-coated dishes. *n*_cell_ = 286 — 297, *n*_exp_ = 2. (**A**) Cumulative distance over time (mean ±s.e.m.). (**B**) Instantaneous speed over time (mean ±s.e.m.). (**C**) Histogram of instantaneous speed. (**D**) Directionality ratio over time (mean ±s.e.m.). (**E**) Mean squared displacement (mean ±s.e.m.). (**F**) Direction autocorrelation (mean ±s.e.m.). (**G**) Histogram of cell turning. The angle distribution for all tracks in the analysis measured from one trajectory to the next. (**I**) Violin plot of average speed of each cell. (**I**) Violin plot of the fastest time taken for each cell to traverse a “segment” of 25 μm in a track. Dots represent individual cells, marker indicates the mean.

**Figure S5.**
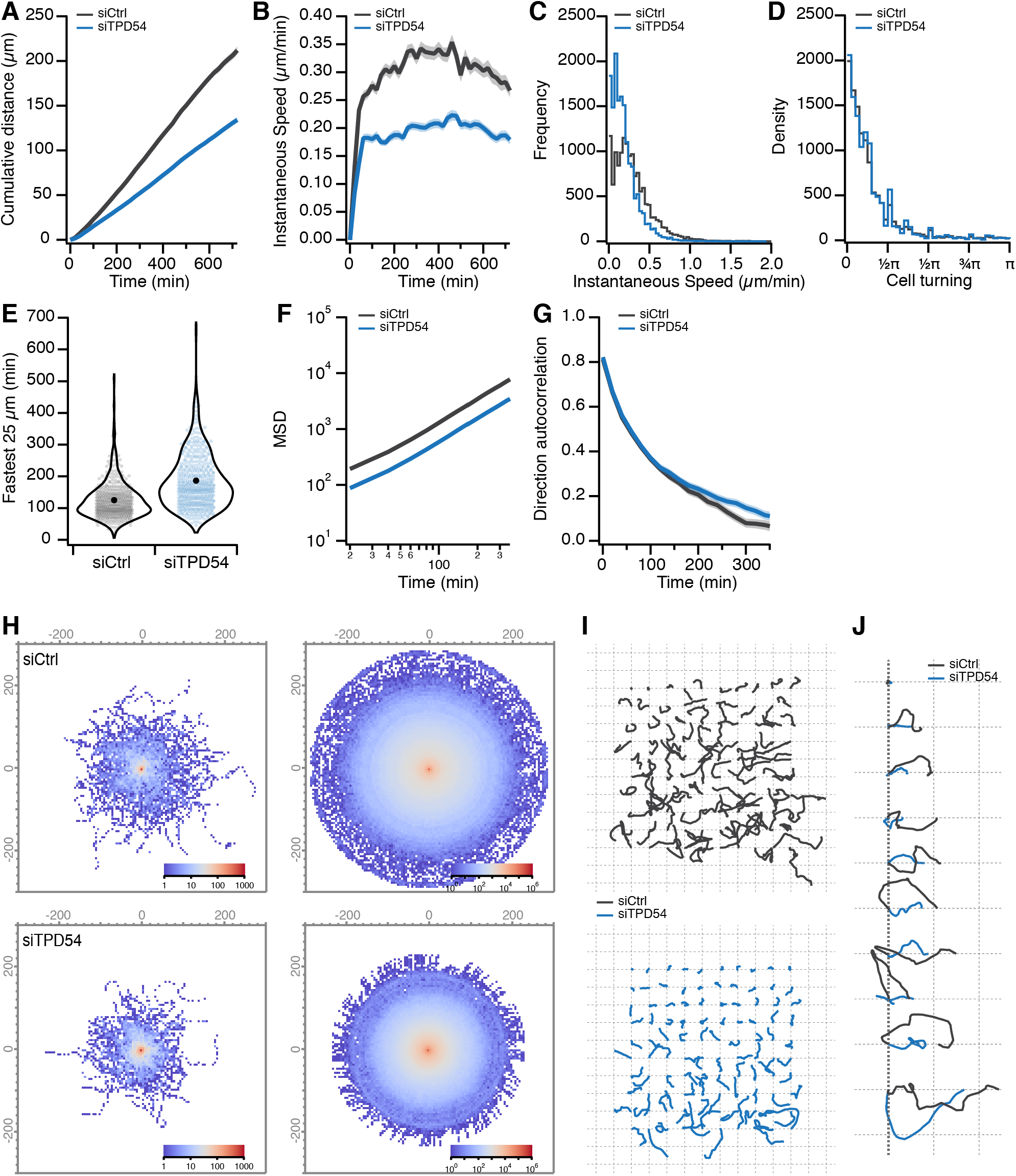
Effect of TPD54 depletion on migration of RPE1 cells. Statistics of Control (gray) or TPD54-depleted (blue) RPE1 cells migrating on fibronectin-coated dishes. Data from Figure 6. (**A**) Cumulative distance over time (mean ±s.e.m.). (**B**) Instantaneous speed over time (mean ±s.e.m.). (**C**) Histogram of instantaneous speed. (**D**) Histogram of cell turning. The angle distribution for all tracks in the analysis measured from one trajectory to the next. (**E**) Violin plot of the fastest time taken for each cell to traverse a “segment” of 25 μm in a track. Dots represent individual cells, marker indicates the mean. (**F**) Mean squared displacement (mean ±s.e.m.). (**G**) Direction autocorrelation (mean ±s.e.m.). (**H**) Overlay of all tracks in the dataset shown as a heatmap (left), bootstrapped + rotated view of cell tracks to visualize the average explored space by cells in the dataset. Density is shown by the log colorscale indicated. (**I**) Image quilt of a sample of cell tracks from the dataset. Each track is shown in its original orientation, arrayed on a grid from the shortest distance traveled to the longest. (**J**) Sparkline image of a diagonal sample through the image quilt (I). To visualize directionality, tracks are rotated such that the end point of the track due east from the start.

**Figure S6.**
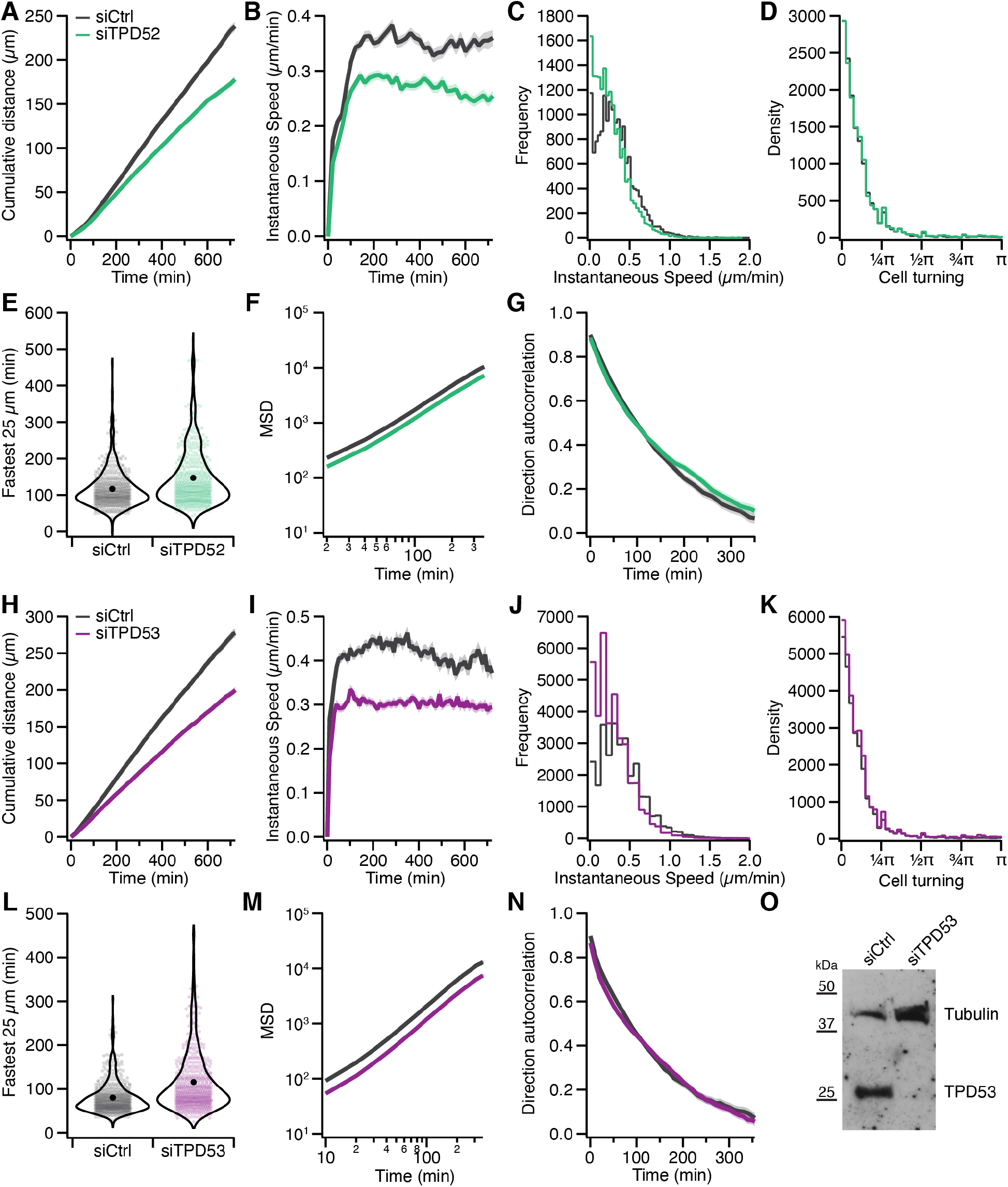
Effect of TPD52 or TPD53 depletion on migration of RPE1 cells. Statistics of Control (gray), TPD52-depleted (green) or TPD53-depleted (purple) RPE1 cells migrating on fibronectin-coated dishes. Data from Figure 6. (**A,H**) Cumulative distance over time (mean ±s.e.m.). (**B,I**) Instantaneous speed over time (mean ±s.e.m.). (**C,J**) Histogram of instantaneous speed. (**D,K**) Histogram of cell turning. The angle distribution for all tracks in the analysis measured from one trajectory to the next. (**E,L**) Violin plot of the fastest time taken for each cell to traverse a “segment” of 25 μm in a track. Dots represent individual cells, marker indicates the mean. (**F,M**) Mean squared displacement (mean ±s.e.m.). (**G,N**) Direction autocorrelation (mean ±s.e.m.). (**O**) Western blotto show depletion of TPD53 under the conditions of the migration experiment. Depletion of TPD52 was not possible to assess due to lack of specific antibodies.

**Figure S7.**
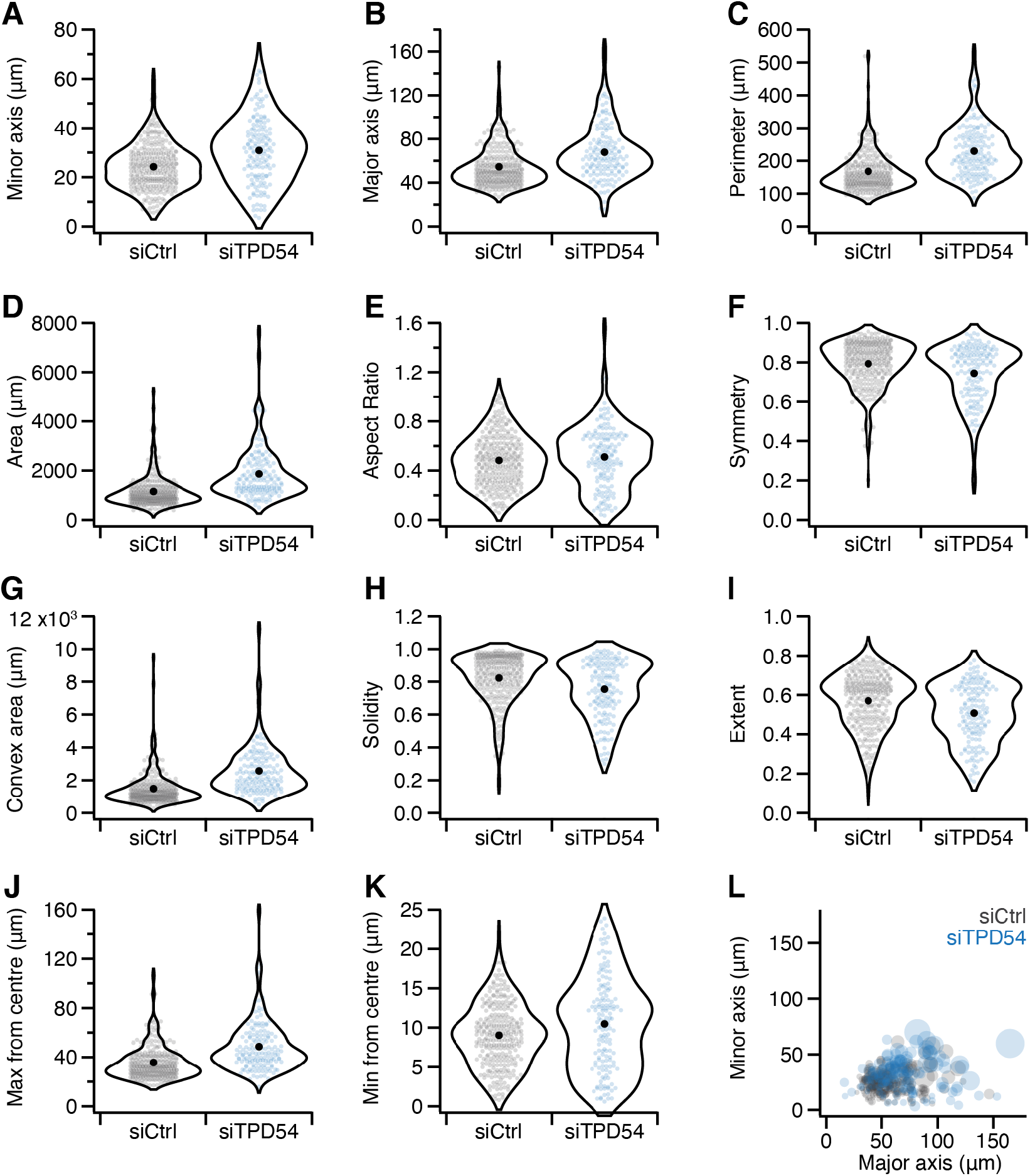
Effect of TPD54 depletion on the shape of RPE1 cells. Cell shape descriptors for control (gray) or TPD54-depleted (blue) RPE1 cells on fibronectin-coated dishes. Valid shapes were analyzed from *n*_cell_ = 288 (siCtrl) and *n*_cell_ = 151 (siTPD54). Violin plots where dots represent individual cells and marker indicates the mean, showing minor axis length (**A**), major axis length (**B**), cell perimeter (**C**), cell area (**D**), aspect ratio, minor:major axes(**E**), symmetry, ratio of the cell area to the area of a the cell footprint reflected on its major axis (**F**), convex area, area of a convex hull enclosing the cell perimeter (**G**), solidity, ratio of cell area to convex area (**H**), extent, ratio of cell area to the area of a bounding box (**I**), maximum distance to the perimeter from the cell center (**J**), minimum distance to the perimeter from the cell center (**K**). (**L**) A plot of minor axis length as a function of major axis length. Each cell in the dataset is represented by a bubble the size of which corresponds to cell area.

## Supplementary Videos

**Figure SV1.**
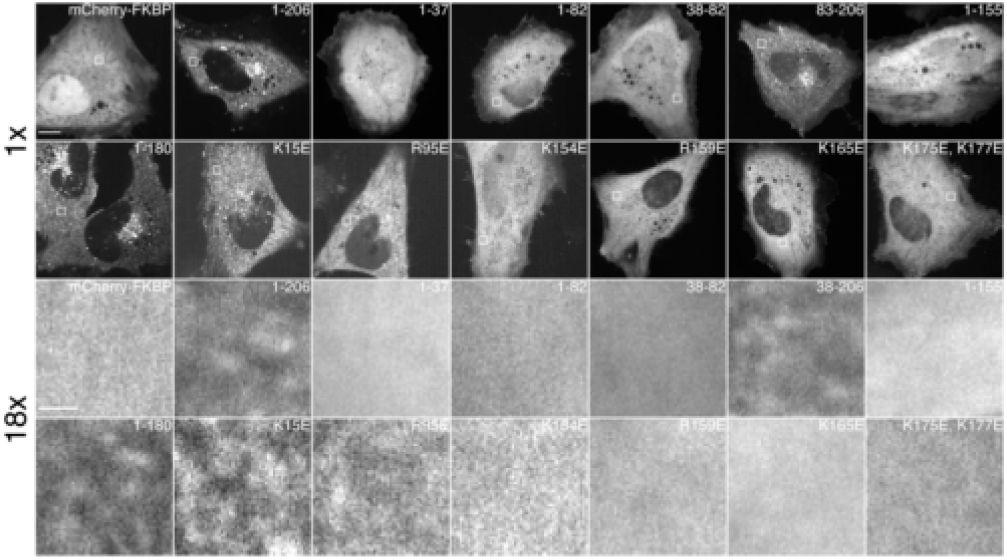
Comparison of fluorescence variance of TPD54 mutants. Representative movies of HeLa cells expressing mCherry-FKBP or mCherry-FKBP-TPD54 constructs, as indicated. Movies were captured at 3.33 Hz. Boxed regions are shown below zoomed to 18×. Scale bar, 10 μm (1 ×) and 1 μm (18×).

**Figure SV2.**
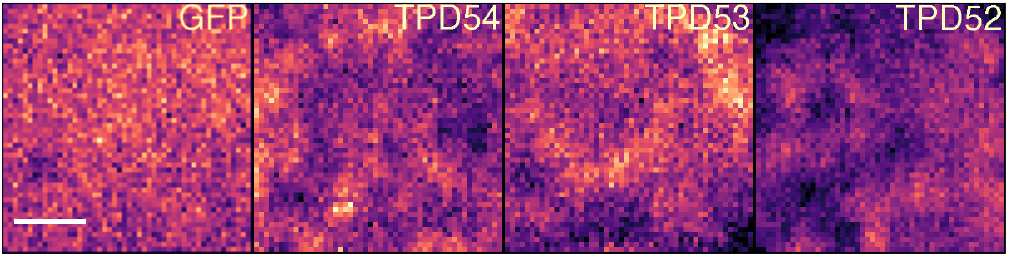
Imaging of TPDs on subresolution vesicles. Live-cell confocal movies of HeLa cells expressing GFP, TPD54, TPD53, or TPD52, captured at ~8 Hz. Scale bar, 1 μm.

**Figure SV3.**
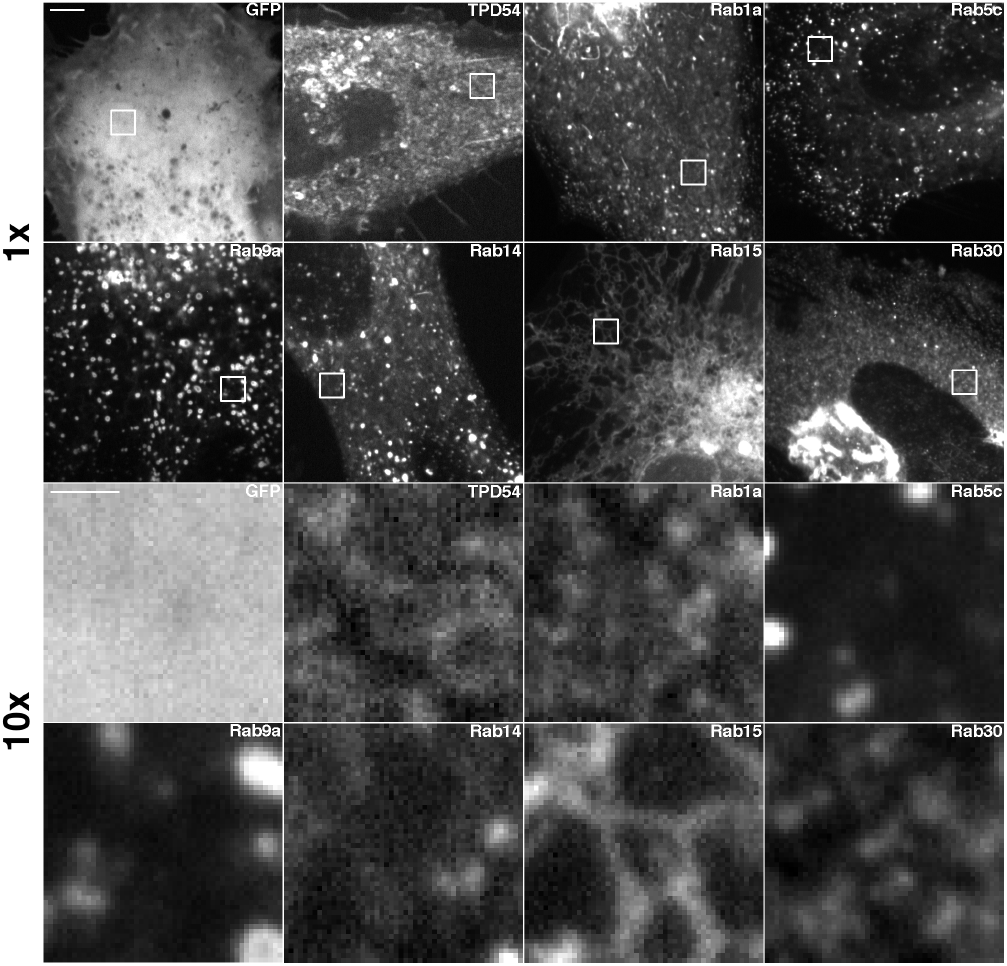
Comparison of fluorescence variance of Rab GTPases. Representative movies of HeLa cells expressing GFP, GFP-TPD54 or GFP-Rabs, captured at 3.33 Hz. Boxed regions are shown below zoomed to 10×. Scale bar, 5μm (1 ×) and 1 μm (10×).

**Figure SV4.**
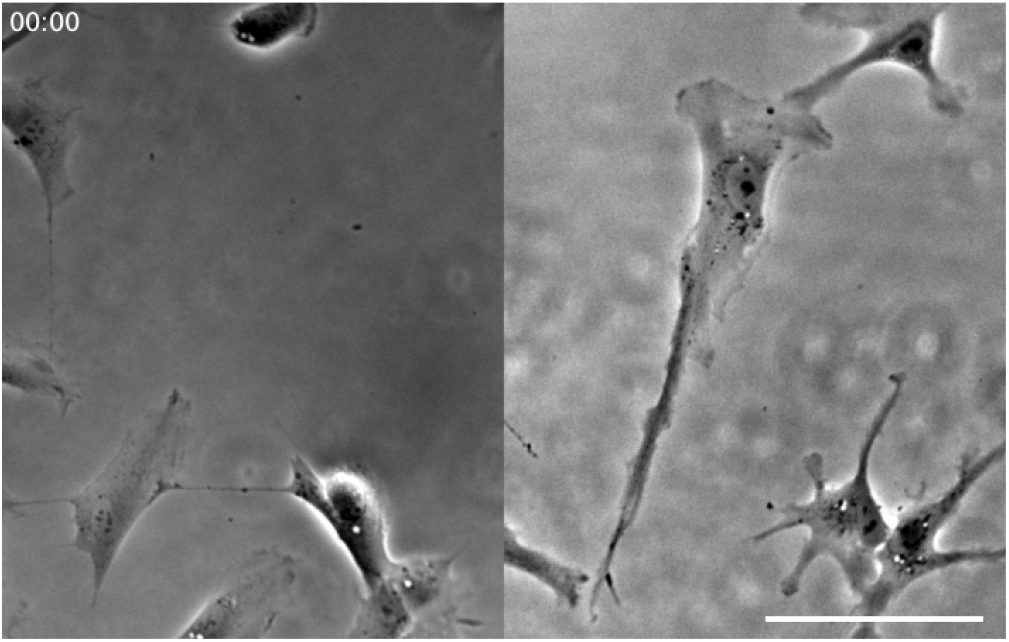
Effect of TPD54 depletion on RPE1 cell morphology and migration. Excerpts of a 12 h recording of RPE1 cells (siControl, left; siTPD54, right) migrating on fibronectin. Scale bar, 100 μm; Time, hh:mm.

## Notes

https://github.com/quantixed/p054p031/

### Competing Interest Statement

The authors have declared no competing interest.

